# The STRIPAK signaling complex regulates phosphorylation of GUL1, an RNA-binding protein that shuttles on endosomes

**DOI:** 10.1101/2020.05.01.072009

**Authors:** V Stein, B Blank-Landeshammer, K Müntjes, R Märker, I Teichert, M Feldbrügge, A Sickmann, U Kück

## Abstract

The striatin-interacting phosphatase and kinase (STRIPAK) multi-subunit signaling complex is highly conserved within eukaryotes. In fungi, STRIPAK controls multicellular development, morphogenesis, pathogenicity, and cell-cell recognition, while in humans, certain diseases are related to this signaling complex. To date, phosphorylation and dephosphorylation targets of STRIPAK are still widely unknown in microbial as well as animal systems. Here, we provide an extended global proteome and phosphoproteome study using the wild type as well as STRIPAK single and double deletion mutants from the filamentous fungus *Sordaria macrospora.* Notably, in the deletion mutants, we identified the differential phosphorylation of 129 proteins, of which 70 phosphorylation sites were previously unknown. Included in the list of STRIPAK targets are eight proteins with RNA recognition motifs (RRMs) including GUL1. Knockout mutants and complemented transformants clearly show that GUL1 affects hyphal growth and sexual development. To assess the role of GUL1 phosphorylation on fungal development, we constructed phospho-mimetic and -deficient mutants of GUL1 residues S180, S216, and S1343. While the S1343 mutants were indistinguishable from wildtype, phospho-deficiency of S180 and S216 resulted in a drastic reduction in hyphal growth and phospho-deficiency of S216 also affects sexual fertility. These results thus suggest that differential phosphorylation of GUL1 regulates developmental processes such as fruiting body maturation and hyphal morphogenesis. Moreover, genetic interaction studies provide strong evidence that GUL1 is not an integral subunit of STRIPAK. Finally, fluorescence microcopy revealed that GUL1 co-localizes with endosomal marker proteins and shuttles on endosomes. Here, we provide a new mechanistic model that explains how STRIPAK-dependent and - independent phosphorylation of GUL1 regulates sexual development and asexual growth.

**Author Summary:** In eukaryotes, the striatin-interacting phosphatase and kinase (STRIPAK) multi-subunit signaling complex controls a variety of developmental processes, and the lack of single STRIPAK subunits is associated with severe developmental defects and diseases. However, in humans, animals, as well as fungal microbes, the phosphorylation and dephosphorylation targets of STRIPAK are still largely unknown. The filamentous fungus *Sordaria macrospora* is a well-established model system used to study the function of STRIPAK, since a collection of STRIPAK mutants is experimentally accessible. We previously established an isobaric tag for relative and absolute quantification (iTRAQ)-based proteomic and phosphoproteomic analysis to identify targets of STRIPAK. Here, we investigate mutants that lack one or two STRIPAK subunits. Our analysis resulted in the identification of 129 putative phosphorylation targets of STRIPAK including GUL1, a homolog of the RNA-binding protein SSD1 from yeast. Using fluorescence microscopy, we demonstrate that GUL1 shuttles on endosomes. We also investigated deletion, phospho-mimetic, and -deletion mutants and revealed that GUL1 regulates sexual and asexual development in a phosphorylation-dependent manner. Collectively, our comprehensive genetic and cellular analysis provides new fundamental insights into the mechanism of how GUL1, as a STRIPAK target, controls multiple cellular functions.

## Introduction

Eleven years ago, an affinity purification/mass spectrometry approach using human cells identified a novel large multiprotein assembly referred to as the striatin-interacting phosphatase and kinase (STRIPAK) multi-subunit complex (1). Besides catalytic (PP2Ac) and scaffolding (PP2AA) subunits of protein phosphatase PP2A, this complex contains regulatory PP2A subunits of the B’’’ family (striatins), which were detected previously in human brain cells as well as in filamentous fungi (2, 3). Further constituents of the core complex include the striatin-interacting proteins STRIP1/2, Mob3/phocein, cerebral cavernous malformation 3 (CCM3), and two associated subunits, the sarcolemmal membrane-associated protein (SLMAP), and the coiled-coil protein suppressor of IκB kinase-ε (IKKε) designated as SIKE (4). STRIPAK is highly conserved within eukaryotes and was shown to control a variety of developmental processes. For example, in filamentous fungi, cell fusion, multicellular fruiting body formation, symbiotic interactions, and pathogenic interactions are dependent on a functional STRIPAK complex. Similarly, several human diseases, such as seizures and strokes, are linked to defective STRIPAK subunits (4-7). Moreover, the phosphorylation activity of STRIPAK is dependent on germinal center kinases (GCKs) such as SmKin3 and SmKin24 (8-10). However, despite major progress in biochemically characterizing STRIPAK complexes, the nature of the upstream regulators and downstream targets affecting signal transduction is not yet fully understood.

To directly address this issue, we have recently performed extensive isobaric tagging for relative and absolute quantification (iTRAQ)-based proteomic and phosphoproteomic analysis to identify putative STRIPAK targets (11). In a wild-type strain from the filamentous fungus *Sordaria macrospora*, we identified 4,193 proteins and 2,489 phosphoproteins, which are represented by 10,635 phosphopeptides (11). By comparing phosphorylation data from wild-type and derived mutants lacking single subunits of STRIPAK, we identified 228 phosphoproteins with differentially regulated phosphorylation sites. Using the iTRAQ quantification method, we compared the relative abundance of phosphorylated peptides in mutants relative to the wild type. Thus, we were able to identify potential dephosphorylation targets of STRIPAK.

Here, we have expanded on our recent analysis by analyzing double mutants. Previous comparative phosphoproteomic studies using double kinase mutants from *Arabidopsis thaliana* showed that phosphorylation states of low-abundance proteins are difficult to detect with either single mutant since background phosphorylation by the other kinase may mask individual targets (12). Similarly, a quantitative phosphoproteomic study with two serine/threonine protein kinases involved in DNA repair revealed that only an *A. thaliana* double mutant enabled kinase-dependent and –independent phosphorylation events to be distinguished between (13). These studies prompted us to expand our recent phosphoproteomic analysis by including double mutants. Here, we analyzed two STRIPAK double mutants lacking either (1) the striatin-interacting protein PRO22 as well as the striatin homolog PRO11 or (2) PRO22 as well as the catalytic PP2A subunit PP2Ac1, with the aim of identifying novel putative targets of STRIPAK, thus increasing the number of potential phosphorylation/dephosphorylation substrates. In this context, we detected GUL1, a homolog of GUL-1 from *Neurospora crassa*, and SSD1 from yeast. In *N. crassa*, the gulliver (*gul*) mutation was identified in a screen for morphological mutants, which act as dominant modifiers of the temperature-sensitive colonial gene *cot*, encoding a NDR kinase. This kinase is a key component of the <u>m</u>orphogenesis <u>or</u>b6 (MOR) network (14-16). Previously, a functional analysis showed that inactivation of the *N. crassa gul-1* gene results in a defect of hyphal polarity as well as cell wall remodeling and hyphal morphology (16, 17). A recent RNAseq analysis with GUL-1 deletion strains identified further genes involved in mycelium development, transcriptional regulation, cell wall biosynthesis, and carbohydrate metabolic processes. Finally, live imaging showed that GUL-1 movement is microtubule-dependent (18). A homolog of GUL1 in yeast is the <u>s</u>uppressor of the <u>S</u>IT4 protein phosphatase <u>d</u>eletion (SSD1), which was discovered in a mutant screen to suppress the lethality of *sit4* deletion strain (19). Later on, SSD1 was shown to be an mRNA-binding protein (20) that shuttles between the nucleus and cytoplasm – a process that is dependent on its phosphorylation state (21-23). Here, we provide a functional analysis of GUL1, which was shown previously in three independent mass spectrometry experiments to associate with the STRIPAK subunit PRO45, a homolog of mammalian SLMAP (24). Our results demonstrate that GUL1 controls sexual development and hyphal morphology in a phosphorylation-dependent manner.

## Results

### iTRAQ-based proteomic and phosphoproteomic analysis of STRIPAK double mutants identifies novel putative targets of the STRIPAK complex

Recently, we have described an iTRAQ-based proteomic and phosphoproteomic analysis of wildtype and mutant strains from *S. macrospora* (11). Here, this analysis was substantially expanded by analyzing two double mutant strains in addition to a single mutant one. In detail, these were Δpro11, lacking striatin, the B’’’ regulatory subunit of phosphatase PP2A, Δpro11Δpro22, lacking striatin as well as PRO22, the homolog of the human striatin-interacting protein STRIP1/2, and Δpp2Ac1Δpro22, lacking PRO22 and the catalytic subunit of PP2A. Protein extracts were isolated from strains grown for 3 days in liquid surface cultures. Two biological replicates were used for each strain. After tryptic digestion, samples were in parallel subjected to global proteome analysis by HPLC-MS/MS and phosphoproteome analysis by TiO_2_ enrichment of phosphopeptides, followed by HILIC, nano-HPLC, and MS/MS as described in Märker et al. (2020). In our analysis, we identified and quantified the global expression levels of a total of 4,349 proteins in all mutant strains and the wildtype, along with the quantification of 9,773 phosphorylated peptides (Fig 1A). The phosphorylation sites in these peptides were localized with high confidence (phosphoRS probability ≥ 90%) and cover a total of 8,177 phosphorylation sites in 2,465 proteins. The expression levels of 1,180 of these phosphoproteins were also determined in the global proteome data, thereby allowing for the differentiation between changes in the phosphorylation level and changes of overall protein expression. The phosphorylation sites showed a distribution of 80 %, 19 %, and 1 % to serine, threonine, and tyrosine residues, respectively. Compared to our recent study (11), we were able to obtain a similar coverage of the proteome, displayed in an overlap of 93 % of all identified proteins (S1A Fig). By using the same deletion strain (Δpro11) in both studies, we were further able to compare the quantitative values with respect to the wild type. Using the log2-ratios of Δpro11 to the wild type of all commonly identified proteins, we calculated a Pearson’s correlation coefficient of 0.73 (S1B Fig). Similarly, the overlap of the phosphoproteomics data amounted to 84% on the level of phosphoproteins and 68 % on the level of phosphosites, with a Pearson’s correlation coefficient of 0.62 for the commonly identified phosphopeptides. Enrichment analysis of the identified upregulated phosphorylation sites showed similar motifs to the ones we identified in the previous analysis (S2A, S2B Fig).

**Fig 1.**
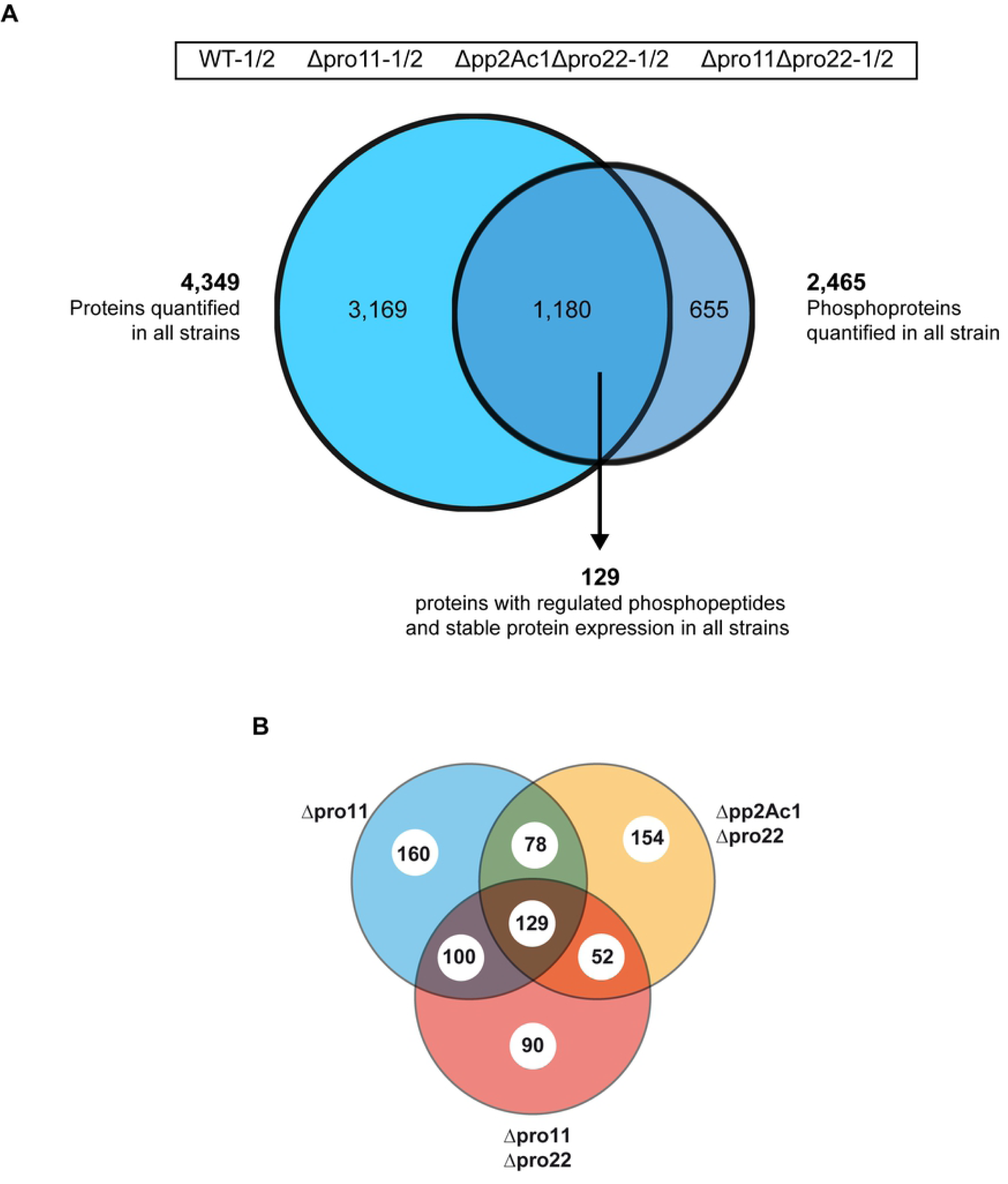
Proteins and phosphoproteins found in the wild type and three STRIPAK deletion strains. (A) We analysed the proteome and the phosphoproteome of the wild type, Δpro11, Δpp2Ac1Δpro22 and Δpro11Δpro22. In total, we identified 4,349 proteins in all strains and 2,465 phosphoproteins. The intersection of the Venn diagram gives the number of proteins found in both analyses (1,180). Moreover, the number of regulated phosphoproteins from all strains are given that were identified with similar abundances in the global proteome. (B) Venn diagram of 129 phosphoproteins with regulated phosphorylation sites in STRIPAK deletion strains. Given is the total number of 129 phosphoproteins in the intersection of the Venn diagram which are differentially phosphorylated in Δpro11, Δpp2Ac1Δpro22, Δpro11Δpro22. Some phosphoproteins are given in more than one intersection because they exhibit multiple regulated phosphorylation sites (see also data sheet S1, S2).

In summary, our analysis revealed a total of 3,624 previously unknown phosphopeptides from 394 previously unknown phosphoproteins (S2C Fig). Further, we identified 342 phosphopeptides to be differentially phosphorylated in all deletion strains of this investigation, and the corresponding 129 proteins were identified in the global proteome without changes in their overall expression levels (Fig 1B, Dataset S1). Of these 129 differentially regulated phosphoproteins, 70 phosphorylation sites were newly identified in this study (11). In Table 1, we provide a selection of newly identified dephosphorylation targets of STRIPAK focusing on those targets that might be involved in signaling during development. Among these targets are HAM5, the scaffold of the pheromone signaling cascade (25, 26), eight kinases, including the STRIPAK-associated GCK SmKIN3 (10), five transcription factors, and eight proteins involved in RNA biology, of which seven contain RNA recognition motifs (27).

**Table 1.**
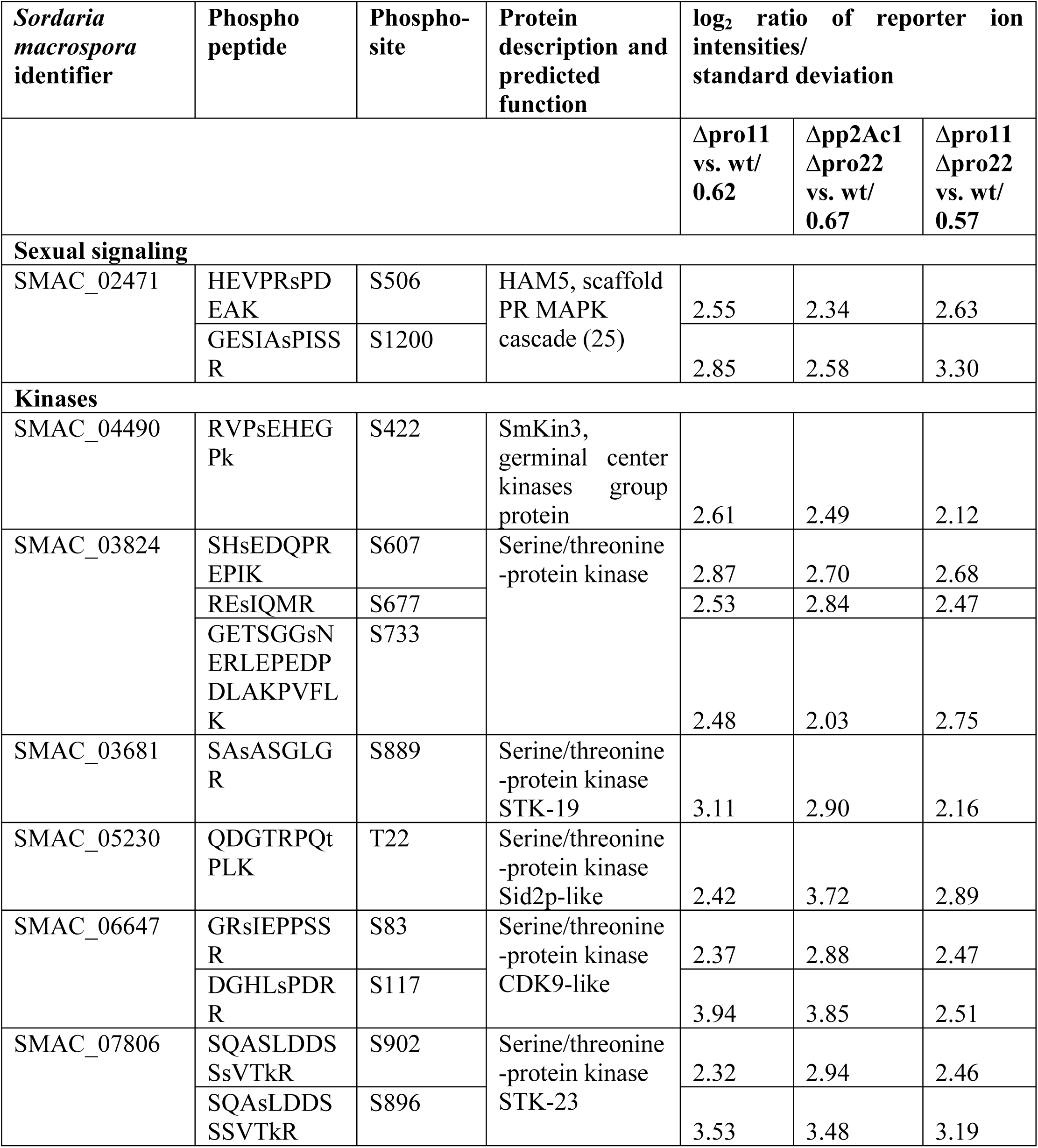
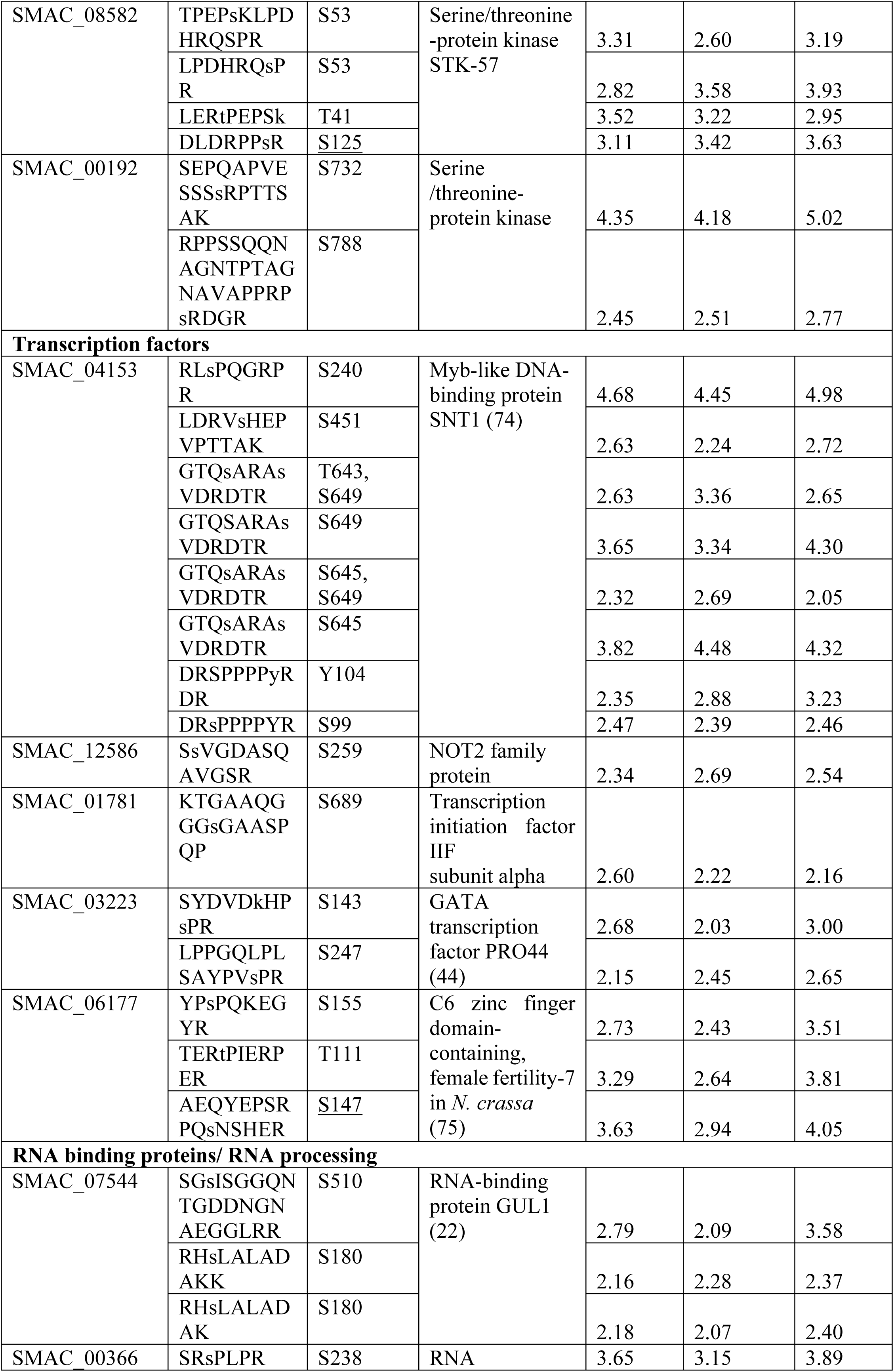
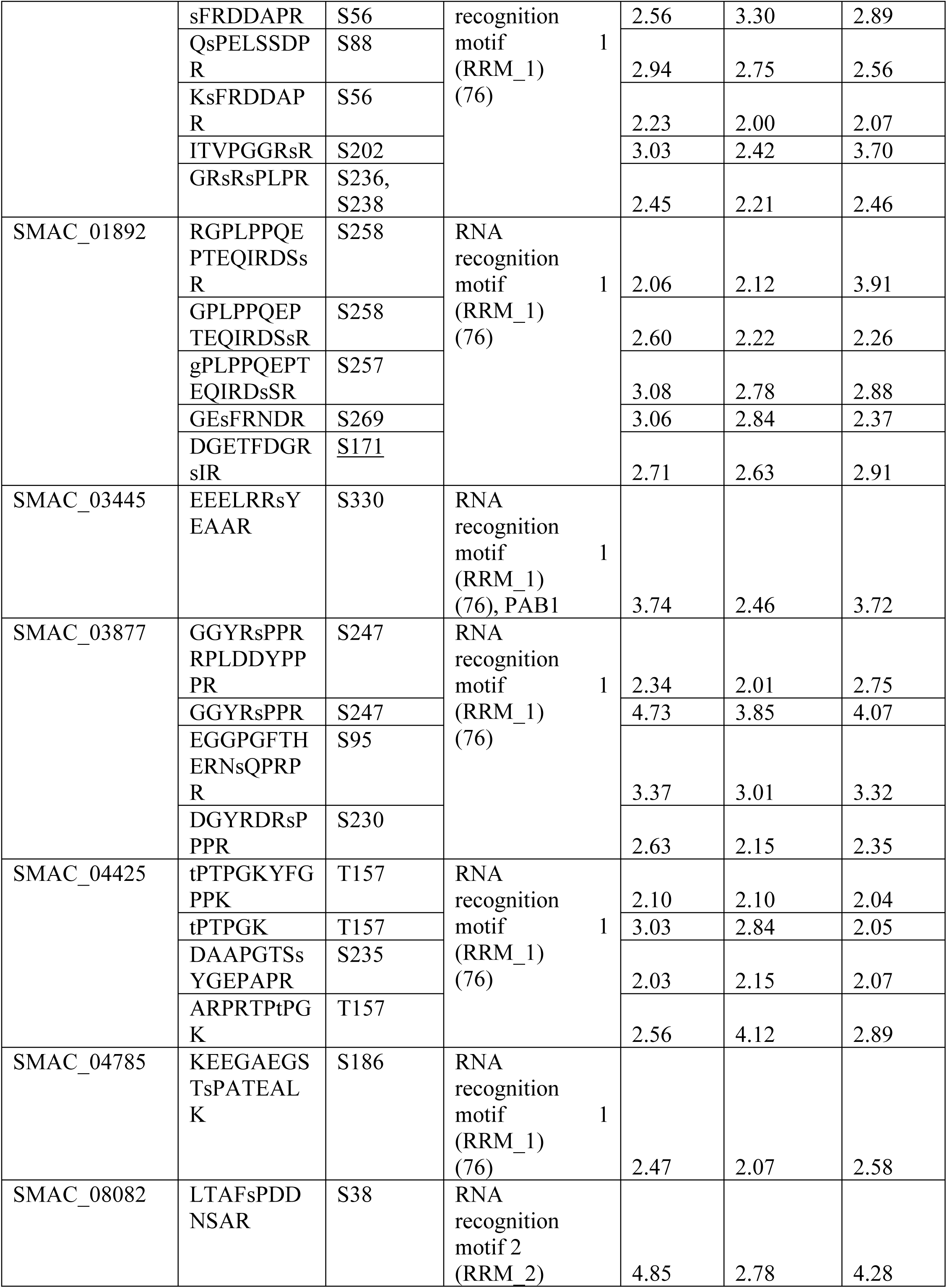
Regulated phosphoproteins in the three investigated STRIPAK mutants Δpro11, Δpp2Ac1Δpro22 and Δpro11Δpro22. Given are 61 differentially phosphorylated peptides from 22 selected proteins. For each phosphorylation site, log2 ratio of reporter ion intensity in deletion strain and wild type relative to the respective standard deviation are given. Underlined are those phosphosites, which were previously detected to be differentially phosphorylated (11). Lower case letters indicate phosphorylated amino acid residues.

Among the 129 putative phosphorylation targets of STRIPAK, we identified 31, which were detected in at least two experiments, as putative interaction partners in previous affinity purification-mass spectrometry analysis with STRIPAK components PRO22, PRO45, or PP2Ac1 as bait (24, 28, 29). For further functional characterization, we chose five out of the 31 newly identified putative targets, namely the putative RhoGAP protein SMAC_06590, the putative nucleoporin SMAC_06564, the catalytic subunit of the mRNA decapping complex SMAC_02163, a ubiquitin-specific protease SMAC_12609, and the RNA-binding protein SMAC_07544, the homolog of GUL-1 from *N. crassa*. Phenotypic analysis of four out of five deletion mutants showed wildtype-like fertility; however, sexual development of Δgul1 was severely affected, as described below.

### GUL1 is a putative dephosphorylation target of the STRIPAK complex

GUL1, carrying an RNA-binding domain, is highly conserved within ascomycetous and basidiomycetous fungi (18, 22, 30). In our phosphoproteomic analysis, we detected ten phosphorylation sites, with phosphorylation of S180 and S510 being differentially regulated in the three mutants of this study (Table 2). Remarkably, differential phosphorylation of S180 was not previously observed in a comparable analysis of three STRIPAK single mutants (11) (S1 Table). This result emphasizes that the investigation of STRIPAK double mutants enables the detection of novel STRIPAK-dependent phosphorylation sites. Further, the finding of numerous RNA-binding proteins suggests that STRIPAK regulates spatio-temporal expression at the posttranscriptional level.

**Table 2.** Identified phosphorylation sites of GUL1. The phosphoproteomic study of Δpro11, Δpp2Ac1Δpro22, and Δpro11Δpro22 compared to the wild type identified ten phosphorylation sites in GUL1. Two out of ten phosphorylation sites are differentially phosphorylated in all three STRIPAK mutants. For each phosphorylation site of GUL1, log2 ratio of reporter ion intensity in deletion strain and wild type relative to the respective standard deviation is given. Bold numbers indicate an upregulation of the phosphorylation site compared to the wild type. Regular numbers indicate no regulation of the phosphorylation site compared to the wild type. The phosphorylation sites marked in red were further analysed in this study (see also Fig 2). Standard deviations: Δpro11: 0.62; Δpp2Ac1Δpro22: 0.67; Δpro11Δpro22: 0.57. In our previous study, we found seven phosphorylation sites (S1 Table.).

|  | Phosphosites |  |  |  |  |  |  |  |  |  |
| --- | --- | --- | --- | --- | --- | --- | --- | --- | --- | --- |
|  | **S180** | S210 | **S216** | S510 | S1198 | T1287 | S1289 | S1291 | T1298 | **S1343** |
| Δpro11/WT | **1.34**<br>**1.35**<br>**1.20** | 0.96 | 0.69 | **1.73**<br>**1.49** | 0.38 | 0.86 | 1.08<br>1.05 | 0.43<br>0.43 | 1.02<br>0.76 | 0.79 |
| Δpp2Ac1Δpro22/WT | **1.53**<br>**1.39**<br>**1.95** | 0.10 | 0.51 | **1.40**<br>1.17 | 0.50 | 0.37 | 0.78<br>0.77 | 0.35<br>0.02 | 1.10<br>0.31 | 0.54 |
| Δpro11Δpro22/WT | **1.35**<br>**1.37**<br>**1.50** | 0.49 | 0.63 | **2.04**<br>**1.26** | 0.74 | **1.65** | 1.30<br>1.00 | 0.63<br>0.24 | 1.02<br>0.54 | 0.59 |

The *gul1* gene carries an open reading frame of 4,353 bp, and encodes for a protein of 1,357 amino acids (27, 31-33). Using the database “eukaryotic linear motif” (ELM) (34), we identified in the primary amino acid sequence of GUL1, a prion-like domain (PLD) (35), several NDR/LATS kinase recognition motifs (36), a nuclear localization signal (NLS), a nuclear export signal (NES), and an RNA-binding domain (Fig 2A). Further, we also detected two consensus sequences for binding of phosphatase PP2A, thus supporting the hypothesis of GUL1 as a target of STRIPAK. This PP2A-binding consensus sequence, as well as the prion-like domain, appear to be absent in the basidiomycetous sequence, while all others are conserved. As shown in Fig 2B, GUL1 shows high sequence identity to homologous proteins in *N. crassa* (NCU_01197; 98 %), *P. anserina* (PODANS_2_6040; 81 %), *M. oryzae* (MGG_08084; 76 %), *F. graminearum* (FG05_07009; 76 %), *S. cerevisiae* (Ssd1p; 42 %), and *U. maydis* (UMAG_01220: 42 %). Our analysis also identified 10 phosphorylation sites, with S180 fitting the NDR/LATS kinase consensus site. Of note is that this region is highly conserved within ascomycetes and basidiomycetes (Fig 2B).

**Fig 2.**
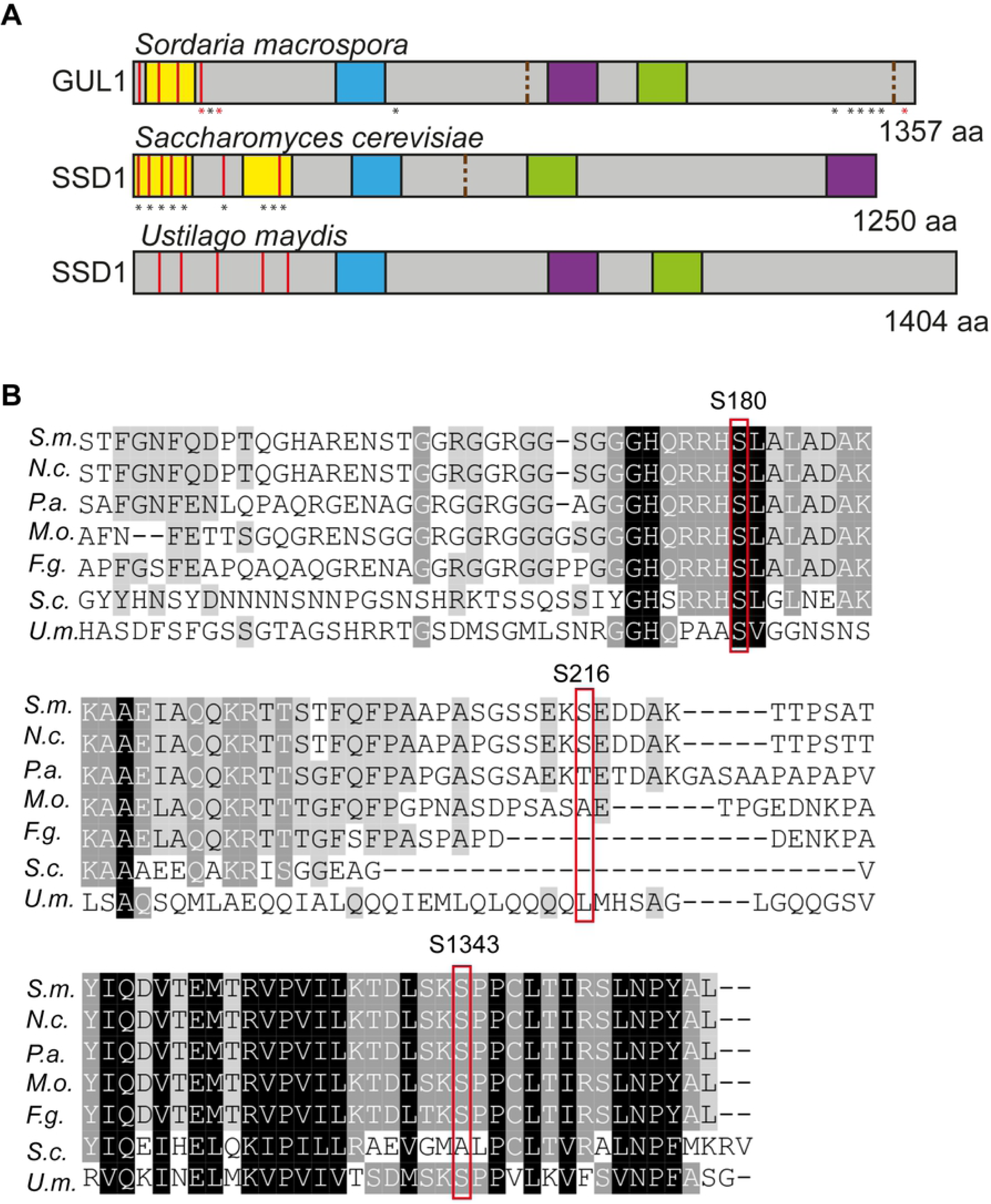
Primary structure and amino acid sequence of GUL1 and its homologues. (A) Identical protein domains in *S. macrospora* GUL1 and its homologue SSD1 in *Saccharomyces cerevisiae* and *Ustilago maydis*. Domains were analysed with ELM and have the following designation: yellow, Prion-like domain; red, LATS/NDR kinase recognition sites; blue, Nuclear localization signal; green, RNA binding domain; purple, nuclear export signal; brown dashed lines, PP2A-binding sites. Asterisks indicate phosphorylation sites, red asteriks in GUL1 were further investigated in this study (S180, S216, S1343). Yeast SSD1 phosphorylation sites were adopted from Kurischko and Broach (2017). (B) Alignment of specific regions of the GUL1 protein from *S. macrospora* sequence with homologues from *N. crassa* (*N.c.*, NCU01197), *P. anserina* (*P.a.*, PODANS_2_6040*), M. oryzae* (*M.o.*, MGG_08084), *F. graminearum* (*F.g.*, FG05_07009), and *S. cerevisiae* (*S.c.*, SCY_1179) and *U. maydis* (*U.m.*, UMAG_01220). Phosphorylation sites S180, S216 and S1343 are framed in red. S180, S216, and S1343 were investigated in the phosphorylation analysis.

### Phosphorylation mutants identify GUL1 residues controlling sexual development and asexual growth

For functional analysis of the *S. macrospora* GUL1 protein, we generated a *gul1* deletion mutant, as described in the Materials and Methods section. The corresponding strain carries a hygromycin B-resistance gene substituting the *gul1* gene by homologous recombination. The deletion strain shows defects in sexual development and in asexual growth (Fig 3). The wildtype forms ascogonial coils, which develop to mature perithecia via protoperithecia within 7 days. *Δgul1* forms all sexual structures including ascogonial coils as well as mature perithecia (Fig 3A). However, the number of ascospores is highly diminished, as can clearly be seen in Fig 3B. Another remarkable phenotype is a defect in hyphal morphology. While hyphae of the wildtype, the complemented strain, and the phospho-mutants are regular and hyphal compartments are straight, hyphae of Δgul1 are hyper-septated and the compartments are swollen (Fig 4A). Compared to hyphal tips, this phenotype is even more severe in mature hyphae (S3 Fig).

**Fig 3.**
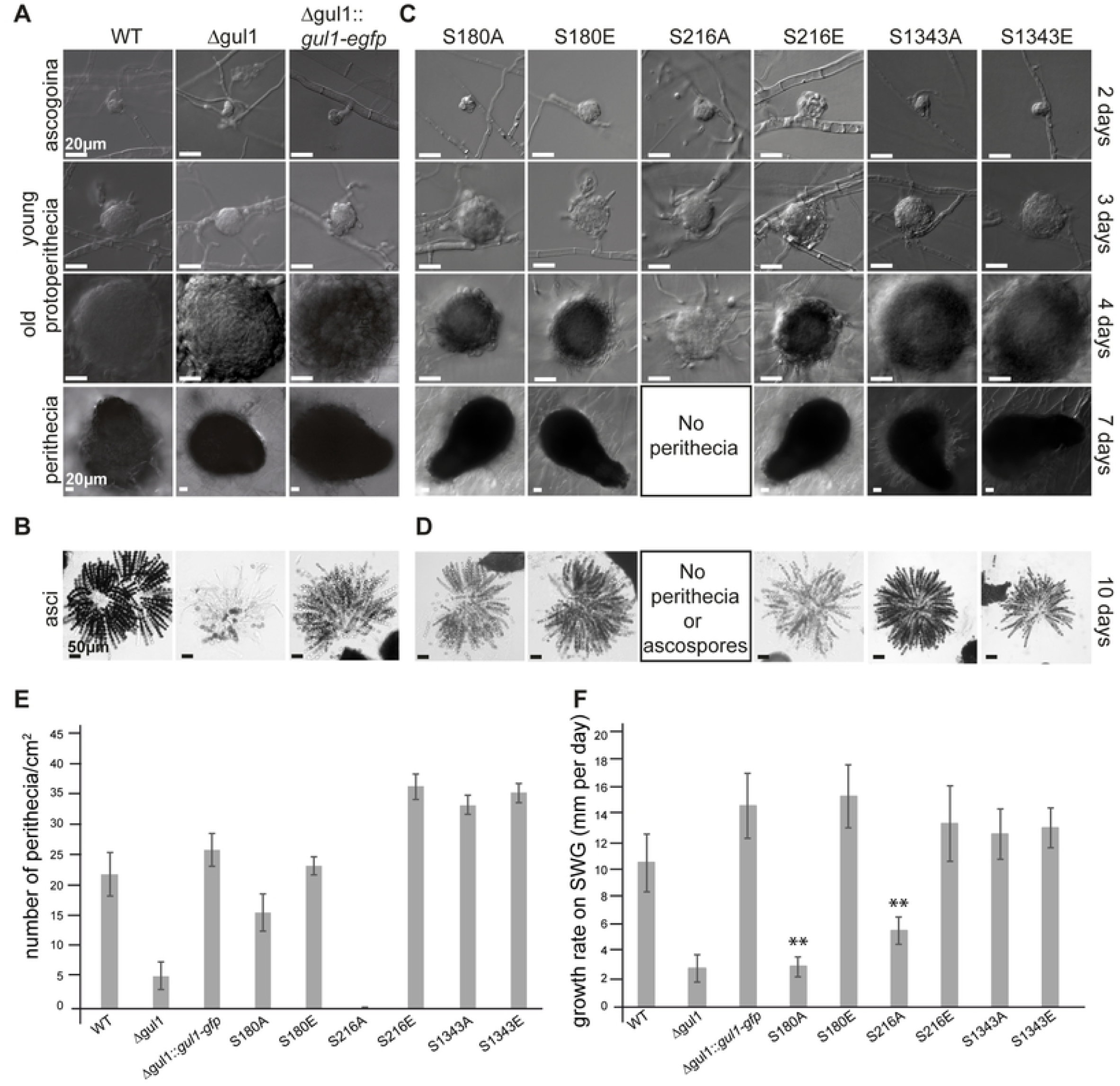
Phenotypic analysis of wild type, Δgul1, a complemented Δgul1 strain (Δgul1::*gul1-gfp*), and phospho-mimetic and – deficient GUL1 strains. (A, B) Sexual development. Ascogonia, young and old protoperithecia, as well as perithecia were examined after 2, 3, 4, and 7 days of growth BMM-slides. Samples were grown on BMM-medium. All bars represent 20 µm. (C, D) Wild type and the Δgul1::*gul1-gfp* complete ascus rosettes, while the *gul*1 deletion strain forms only a few ascospores. Phospho-deficient GUL1 strain S180A and both phospho-mimetic GUL1 strains S180E and S216E show complete ascus rosettes. Phospho-deficient GUL1 strain S216A does not form any spores. Phospho-deficient GUL1 strain S1343A and phospho-mimetic GUL1 strains S1343E show a wild-type like fertility. Bar represent 50 µm. (E) Quantification of perithecia per square centimetre on solid BMM-medium after 10 days (*n* = 9). (F) Growth rate of GUL1 phospho-mutants compared to Δgul1::*gul1-gfp* on SWG. Asterisks indicate significant differences compared to the complemented strain. Error bars in E and F indicate the standard deviation.

**Fig 4.**
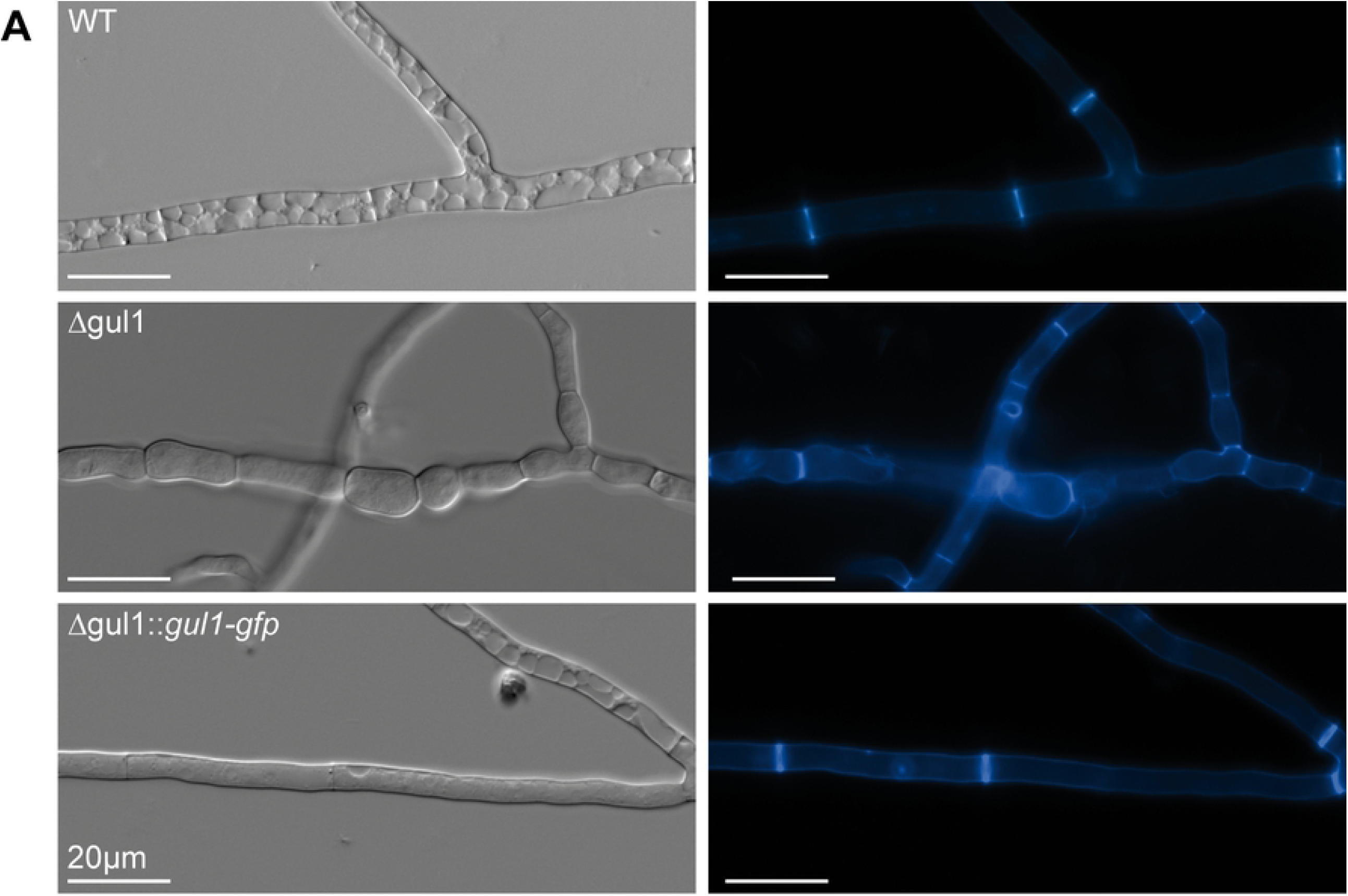
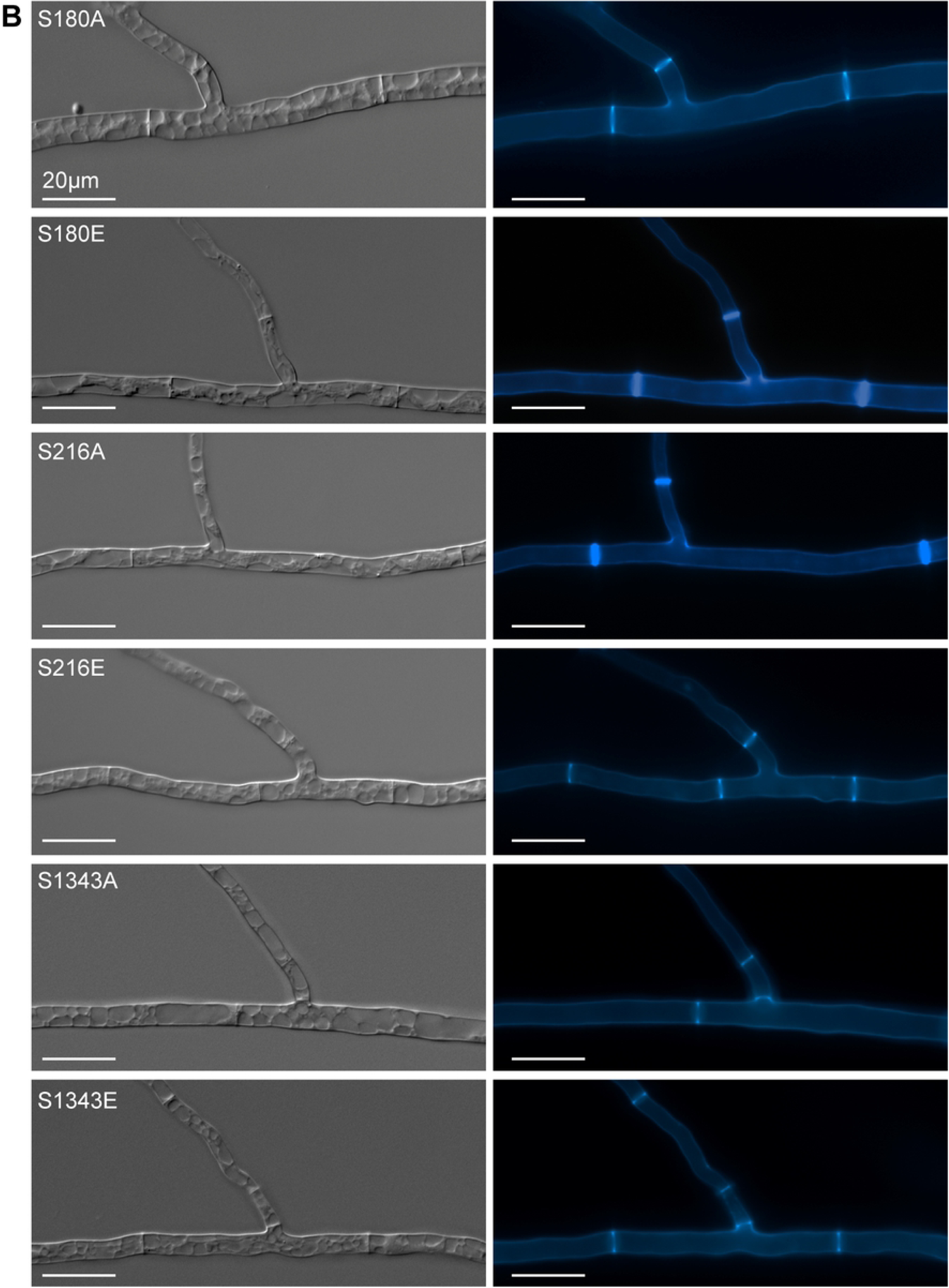
Septation and hyphal morphology of the wt, Δgul1 and the complemented Δgul1-strain Δgul1::*gul1-gfp* compared to phospho-mimetic and – deficient gul1 strains. (A) The septation of hyphae in the wild type as reference, as well as in the complemented strain is regular and hyphal compartments are straight. In the *gul1* deletion strain hyphae are hyperseptated and the compartments appear in a bubble-like structure. (B) Phospho-mimetic and – deficient GUL1 strains show no difference compared to wild type. Strains were grown on MMS for 2 days and stained with Calcofluor White M2R. Bars: 20 µm.

As shown above, we identified GUL1 as a putative dephosphorylation target of STRIPAK. To assess the physiological relevance of GUL1 phosphorylation, we chose three phosphorylation sites for functional analysis. S180, which is STRIPAK dependently phosphorylated (Table 1), is part of a predicted NDR/LATS kinase recognition motif, a highly conserved sequence in all eukaryotes. S216, a less conserved site, is not STRIPAK dependently phosphorylated (Table 1). From the domain analysis, we predict that this site is probably a target of a casein kinase.

Finally, S1343 is C-terminally located in a highly conserved region. As described in the Material and Methods section, the three triplets encoding S180, S216, and S1343 were individually subjected to *in vitro* mutagenesis, resulting in substitution of the corresponding serine triplets to either alanine (prevents phosphorylation) or glutamic acid triplets (mimics phosphorylation because of the negative charge). After transformation of the *gul1* deletion strain with the mutated genes, we investigated three homokaryotic ascospore isolates of each, the phospho-mimetic strains S180E, S216E, and S1343E, the phospho-deficient strains S216A and S1343A, as well as three independent primary transformants S180A (see the Material and Methods sections for construction). Western blot analysis using an anti-GFP antibody detected the corresponding GUL1-GFP fusion proteins and thus confirmed the translational expression of the mutated genes (S4 Fig). All strains were phenotypically characterized concerning fruiting body and ascospore formation as well as vegetative growth (Fig 3). Phospho-deficient and phospho-mimetic strains S1343A and S1343E had similar characteristics compared to wild type (Fig 3C, 3D, 3E, 3F). S180E and S180A were fully fertile generating mature fruiting bodies and ascospores (Fig 3C and 3D); however, the number of perithecia per square centimeter in S180A was considerably reduced by about 25 % compared to wild type (Fig 3E). Further, S180A also showed a reduced growth rate comparable to Δgul1 (Fig 3F). An intriguing result was obtained with S216. While S216E has a wild type phenotype, phospho-deficient strain S216A is sterile and forms only protoperithecia. S216A also has a reduced growth rate comparable to S180A and Δgul1. Thus, this phosphorylation site, which seems not to be targeted by STRIPAK, regulates both sexual and hyphal development (Fig 3C, 3D). Interestingly, none of the six phosphorylation mutants exhibits the severe hyphal swelling phenotype observed in Δgul1 (Fig 4B). In conclusion, we hypothesize that the STRIPAK-dependent phosphorylation of S180 is a switch for hyphal growth, and to some extent, also effects sexual development. In contrast, phosphorylation of S216 is STRIPAK independent, but essential for the formation of mature fruiting bodies as well as hyphal growth. The phosphorylation of S1343 seems to be not essential for sexual development and asexual growth.

### GUL1 is not an integral subunit of the STRIPAK complex

As mentioned above, our previous affinity-purification MS analysis indicated that GUL1 interacts with the STRIPAK subunit PRO45, a homolog of mammalian SLMAP (24). Similarly, SSD1 interacts in a two-hybrid analysis with the yeast protein FAR10, a homolog of PRO45 (37). In addition, this study considered a negative genetic interaction (GI) between *far10* and *ssd1*. Therefore, to determine whether GUL1 is an integral part of the STRIPAK complex or only associated with it, we examined the GI by investigating the double mutant Δpro45Δgul1. For this purpose, we compared the phenotype of the double mutant with the phenotype of the corresponding single mutants by measuring the vegetative growth rates. Compared to wild type, both single and double mutants showed reduced growth rates (Fig 5). We thus used this phenotypical trait to calculate the GI of *gul1* with *pro45.* It is assumed that the phenotype of a double mutant is the result of the phenotype of both single mutants. Whereas a negative GI denotes a reduced fitness of the double mutant compared to both single mutants, a positive GI refers to a higher fitness than expected. Genes encoding for proteins of different pathways often show a negative GI and those encoding for proteins of the same pathway mostly have a positive GI (10, 38, 39). As control, we used the double mutant Δpro45Δpro11 and both single mutants since both are known STRIPAK core subunits and show direct physical interaction (24). Thus, both genes can be considered to have a positive GI. The absolute values of the vegetative growth rates were calculated relative to wild type, with a value of 1 (S2 Table). The data of the single mutants Δpro45, Δpro11, and Δgul1 are 0.494±0.03, 0.413±0.02, and 0.189±0.01, respectively. The expected values were calculated as described previously (10) and are as follows: Δpro45Δpro11, 0.204 and Δpro45Δgul1, 0.093 (see S2 Table). These expected values (light blue bars in Fig 5) were compared to the experimentally obtained values. As expected, the double mutant Δpro45Δpro11 showed no significant deviation of the experimental value from the expected value, indicating the positive GI of *pro11* and *pro45*, as expected. In contrast, the experimentally obtained value for the double mutant Δpro45Δgul1 was significantly lower than the expected values (Fig 5).

**Fig 5.**
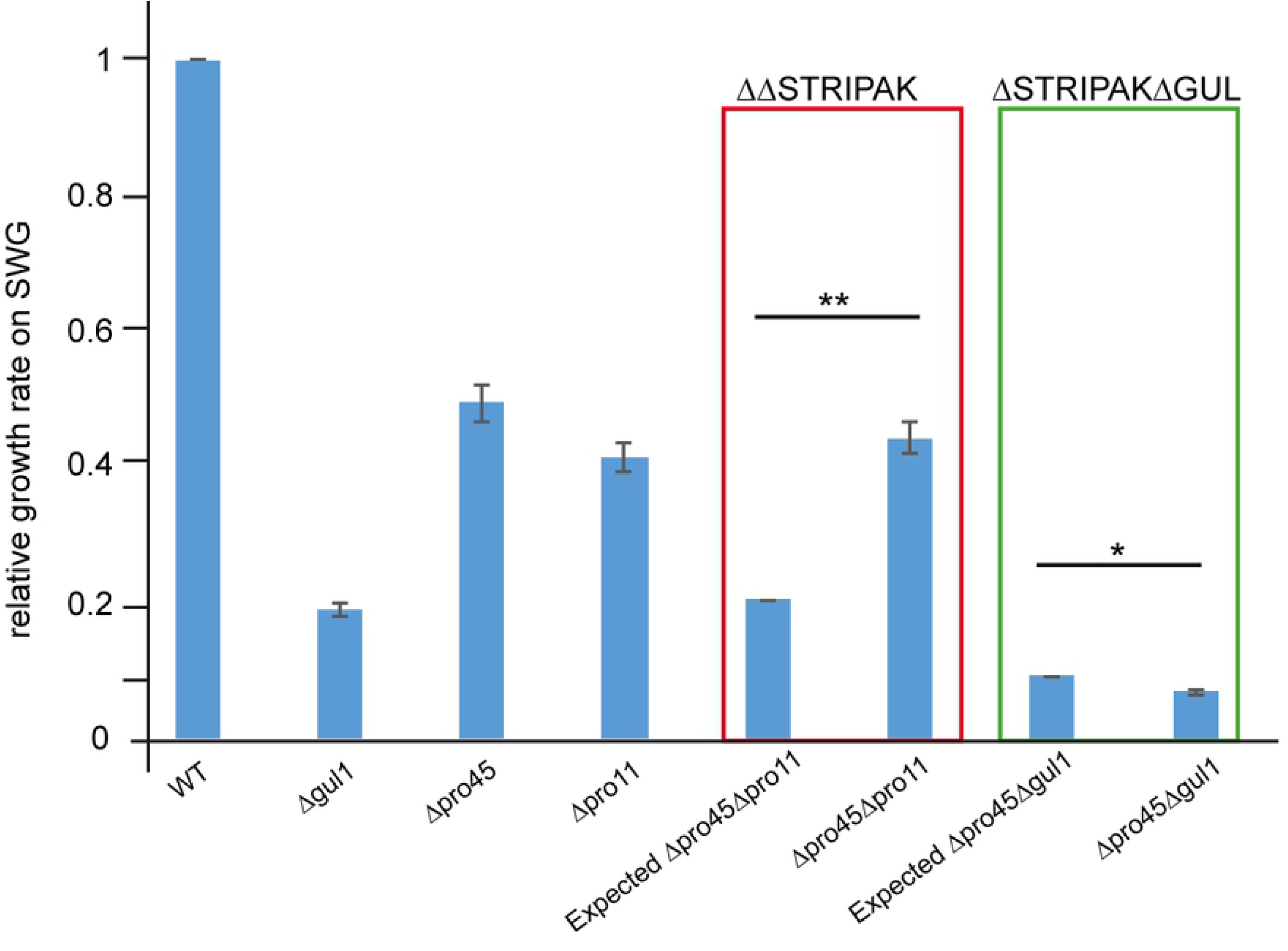
Analysis of the genetic interaction between *gul1* and the *slmap* homologue *pro45*. Genetic interaction was evaluated by comparing the daily vegetative growth rates of the indicated strains. The evaluation is based on the phenotype of the double mutant Δgul1/Δpro45 compared to the single mutants. The double mutant Δpro45/Δpro11 served as a control. Dark blue bars indicate experimentally generated values for single and double mutants, while light blue bars represent expected values for the double mutants based on multiplication of the values of the single mutants. The value of the wild type (wt) was set to 1 and all other values are given in relation to the wt. Absolute and relative values can be found in S2 Table. Error bars indicate standard deviations. Significant differences were evaluated by paired one-tailed Student’s *t-*test and are shown by lines * *P* ≤ 0.05; ** *P* ≤ 0.01. (*n* = 9 see Strains and growth conditions for details).

### GUL1 locates close to the nucleus and shuttles on endosomes

To study the subcellular localization of GUL1 *in vivo*, we analyzed the complemented Δgul1 strain expressing a GUL1-GFP fusion protein. As shown above, this strain shows a wild type-like phenotype, proving the functionality of the fusion protein. Fluorescence microscopy revealed that GUL1 appeared within particle-like structures. These were evenly distributed within the cytoplasm, and some appeared close to nuclei. This observation was further verified when we investigated a strain that expresses both genes for *gul1-gfp* and *h2a-mrfp* (Fig 6). As indicated by red arrows, GUL1 localizes close to nuclei, thereby suggesting a localization to spindle pole bodies.

**Fig 6.**
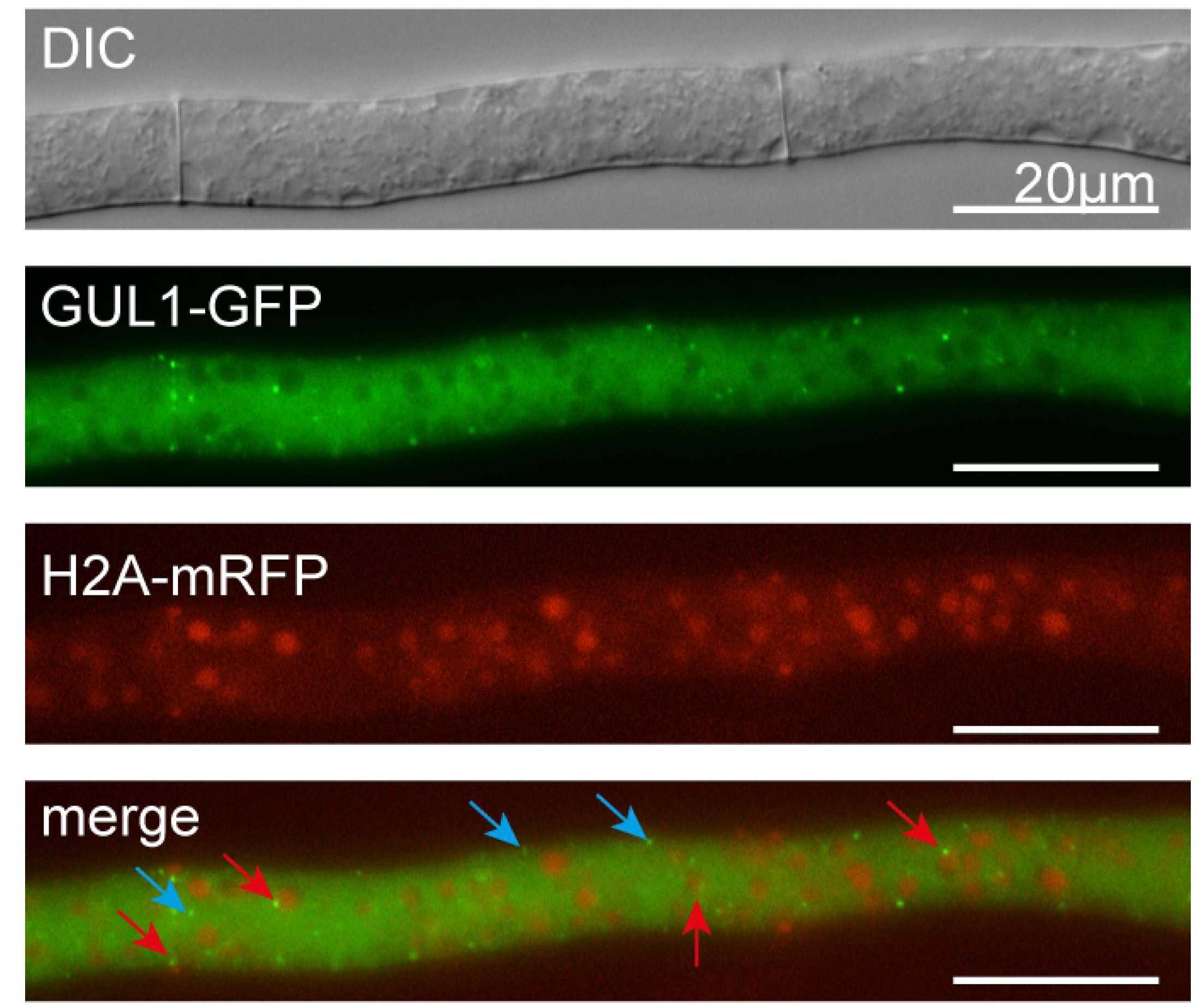
Localization of GUL1-GFP and H2A-mRFP in hyphae of Δgul1. GUL1 localizes in dot-like structures within the cytoplasm (blue arrows). Red arrows indicate a localization of GUL1 close to the nucleus.

To address potential microtubule-dependent movement of GUL1 (18), we performed dynamic live cell imaging (S1 movie). We asked whether the mutation of phosphorylation sites has an effect on long distance movement of GUL1. Analyzing GUL1-GFP expressing strains revealed extensive bidirectional movement of GUL1-GFP, which was most prominent in the vicinity of growing hypha (Fig 7A). The velocity of processive particles was 2.4 µm/s (Fig 7B). We did not observe significant differences analysing GUL1-GFP velocity in the phospho variants (Fig 7B, S5 Fig).

**Fig 7.**
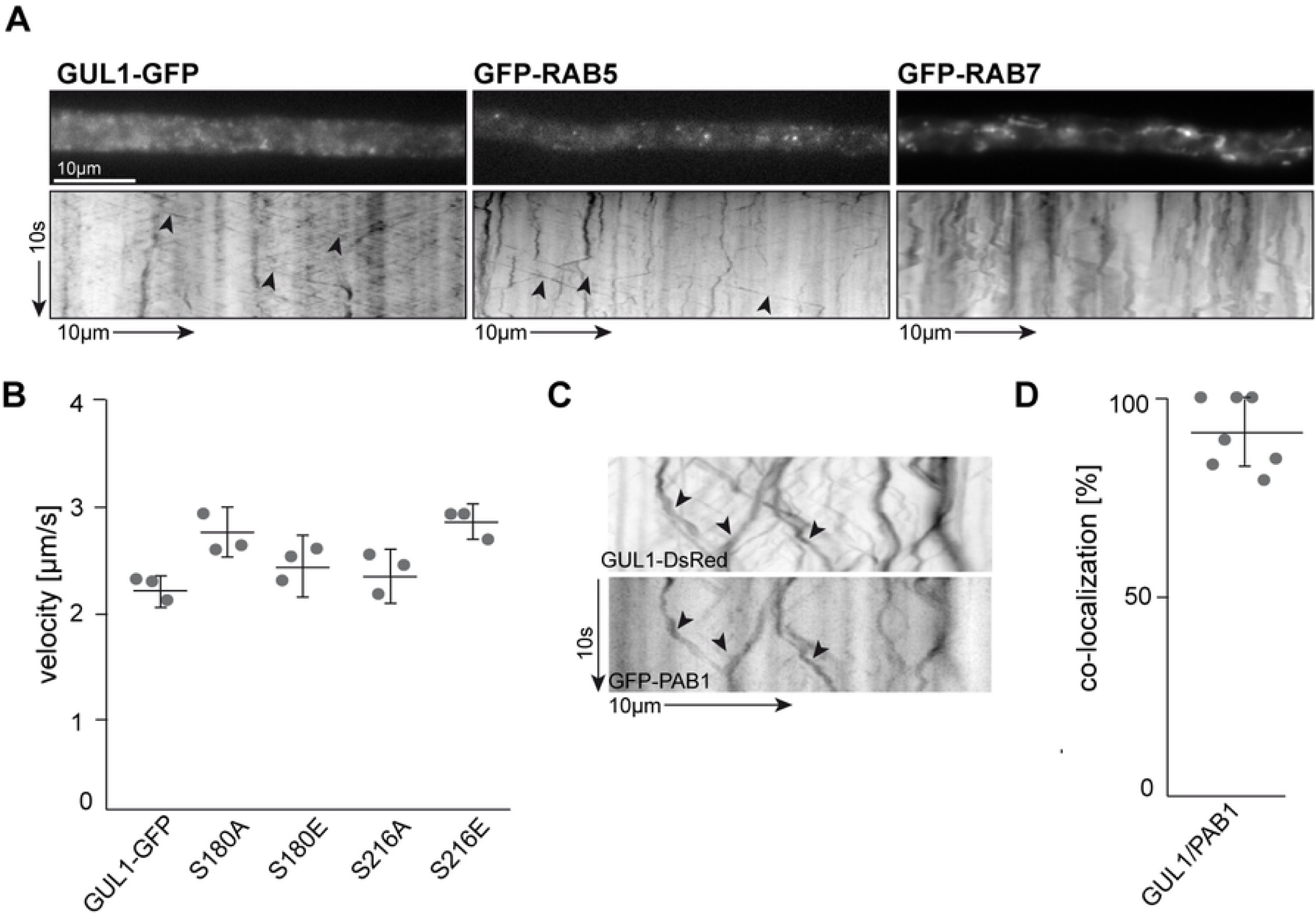
GUL1 co-localizes with PAB1 and shuttles similar to RAB5-positive endosomes throughout hyphae. (A) Kymographs comparing hyphae expressing GUL1-GFP in the *gul1* deletion strain, GFP-RAB5 and GFP-RAB7 in the wild type. Processive signals are marked by black arrowheads; arrow length on the left and bottom indicates time and distance, 10 s and 10 µm, respectively; S1-S3 movie). (B) Average velocity of fluorescent signals per kymograph for strains as indicated. Data points represent three means out of 20 independent hyphae. At least 10 signals/hypha were analysed. Mean is indicated by a black line. (C) Kymograph of a hyphae expressing GUL1-DsRed and GFP-PAB1. Fluorescence signals were detected simultaneously using dual-view technology. Processive co-localizing signals are marked by black arrowheads (S4 movie). (D) Percentage of red fluorescent signals exhibiting co-localization with the green fluorescent signal for strains shown in (C). Data points represent observed co-localization of three means out of seven independent hyphae. Mean is indicated by a horizontal line. Error bars in B and D indicate standard deviation.

Of note, the GUL1-GFP movement is reminiscent of endosomal shuttling in fungi (40, 41). To address this point we studied strains expressing GFP-RAB5 and GFP-RAB7, which are established markers for early and late endosomes. Interestingly, the bidirectional movement of GUL1-GFP resembled the bidirectional shuttling of GFP-RAB5-positive endosomes (Fig 7A, S2 movie). To address a potential role of the RBP GUL1 in endosomal mRNA transport we studied co-localization of GUL1-DsRed with the poly(A) binding protein PAB1 (SMAC_03445) fused to GFP. Importantly, the latter was also identified in our differential phosphorylation study (Table 1). We observed extensive co-localization in processively moving units, suggesting that the RNA-binding protein GUL1 participates in endosomal mRNA transport (Fig 7C-D; S4 movie). Importantly, this is the first evidence that this mode of long-distance transport is also present in ascomycetes (42). Taken together, our fluorescent microscopic investigation reveals that GUL1 acts close to nuclei and shuttles with PAB1 and transport endosomes along microtubules.

## Discussion

The STRIPAK multi-subunit complex is highly conserved within eukaryotes and the number of reports is increasing that single subunits control a huge variety of developmental processes. Despite the intense interest in STRIPAK, our current knowledge about dephosphorylation targets is quite limited and our understanding of how STRIPAK regulates cell differentiation remains basic. Thus, this study provides new fundamental insights into this research field.

We used a quantitative proteomic and phosphoproteomic analysis to identify targets of STRIPAK in the model fungus *Sordaria macrospora*, for which a collection of STRIPAK single and double mutants are available (43). Compared with our recent study (11), we have now gone beyond this by identifying numerous novel STRIPAK dephosphorylation targets. In detail, we identified five transcription factors, such as the GATA transcription factor PRO44, which was shown to control fungal sexual fertility (44). In PRO44, we detected three phosphorylation sites, two of which are differentially regulated in the double mutants. Notably, another protein (SMAC_08582) shows similarity to serine/threonine kinase STK-57 in *N. crassa* (45), and carries four phosphorylation sites of which three are differentially phosphorylated in all STRIPAK mutants investigated in this study. Among these, S125 is also differentially regulated in three single mutants of our recent investigation (11). Another remarkable putative STRIPAK target is HAM5, the scaffold protein of the MAK-2 pathway (25, 26), with 18 phosphorylation sites. Two sites seem to be differentially regulated in single mutants, namely S506 in Δpro11 and Δpro22 as well as S1200 in Δpro22 (11). Interestingly, we also found the differential regulation of both sites in all three STRIPAK mutants. Our investigation of two STRIPAK double mutants detected differentially phosphorylated proteins, which seem to be unique in this experimental approach. For example, we detected the serine/threonine kinase SMAC_00192, which has nine phosphorylation sites, with two (S782, S788) that are differentially regulated. Intriguingly, we also identified numerous potential RNA-binding proteins as targets of STRIPAK, thus suggesting extensive regulation of gene expression by STRIPAK at the posttranscriptional level. Among the candidates were PAB1 (SMAC_03445), a poly(A)-binding protein that shuttles on endosomes (46), as well as GUL1, a regulator of fungal morphogenesis (14, 17).

### GUL1 is involved in different developmental processes

GUL1 is a highly conserved protein in yeast and filamentous fungi, but its cellular function is currently only partly understood. Our analysis has now revealed an RNA-binding domain, a nuclear localization signal, and a nuclear export signal – among others – in the primary sequence of GUL1. These domains have led us to the conclusion that *S. macrospora* GUL1 is an RNA-binding protein, as was previously shown by functional analysis in other filamentous fungi and yeasts (16, 22, 47-49). In the human pathogenic yeast *Candida albicans*, SSD1, the GUL1 homolog, was described as an mRNA-binding protein acting as a translational repressor (49), and the GUL1 homologs in *Magnaporthe oryzae* and *Aspergillus fumigatus* were described as cell wall biogenesis proteins (47, 48). In this study, we provide a comprehensive overview of GUL1’s possible roles, which are related to sexual development, hyphal morphology, as well as vegetative growth. While the *gul1* deletion strain shows a severe reduction in fertility, the phospho-deficient GUL1^S216A^ variant displays a sterile phenotype and both phospho-deficient variants, GUL1^S216A^ and GUL1^S180A^, exhibit severely reduced vegetative growth. Moreover, the sterile phenotype observed in GUL1 and STRIPAK mutants suggests a further association between both, as was previously demonstrated with the STRIPAK-associated GCK SmKIN3 (10). This association, however, is only fully functional if the phosphorylation states of STRIPAK targets are tightly regulated. In essence, we provide compelling evidence that the STRIPAK target GUL1 is extensively regulated at the level of phosphorylation.

### GUL1 is trafficking on endosomes

Fluorescence microcopy showed that GUL1 localizes not only to cytoplasmic punctae, but also close to nuclei, thereby suggesting localization at the nuclear membrane. This hypothesis is further supported by the interaction of GUL1 with the SLMAP homolog, PRO45, which localizes to the nuclear membrane in wild type strains. However, lack of PRO11 or PRO22 is known to prevent nuclear membrane localization of PRO45 (24), which in turn probably reduces the level of dephosphorylation of GUL1. These observations are consistent with data for the GUL1 homolog from yeast. In this case, nucleocytoplasmic shuttling of SSD1 is essential for mRNA binding (21).

Our imaging data provide compelling evidence that in fungal cells GUL1is present on RAB5-positive transport endosomes, which shuttle along microtubules. Consistently, microtubule-dependent movement has been already described for GUL-1 from *N. crassa* (18, 50). Endosomal mRNA transport is well-studied in the basidiomycete *Ustilago maydis* and key components are the RNA-bindings proteins (RBPs) Rrm4, the poly(A)-binding protein PAB1 and the small glycine rich RRM protein Grp1 (40, 51). These RBPs form higher-order transport mRNPs that contain cargo mRNAs encoding e.g. septins for endosomal assembly (51-53). Transport mRNPs are stabilized by the scaffold protein Upa2 and linked to endosomes via the FYVE domain protein Upa1 (41, 46). A phylogenetic analysis revealed that important core components of endosomal transport like Upa2 and the key RBP Rrm4 are missing in ascomycetes (42). However, here we demonstrate that numerous RNA-binding proteins containing RRM domains are prominent STRIPAK targets. Intriguingly, this includes important RBPs of fungal endosomal mRNA transport machinery: the Ssd1 homologue GUL1, PAB1 and a small Glycin rich RRM protein (SMAC_04425) suggesting that STRIPAK regulates this mode of RNA transport. Consistently, GUL1 and PAB1 co-shuttle similar to RAB5-positive endosomes in growing hypha. In essence, we provide compelling evidence that STRIPAK is a posttranscriptional regulator most likely orchestrating endosomal mRNA transport and that this transport mechanism is conserved in all fungi including ascomycetes.

However, we presume that the composition of the endosomal transport complex is slightly different in asco- and basidiomycetes. Importantly, endosomal mRNA transport and local translation on the cytoplasmic surface of endosomes was recently described in neurons (54) and in this context it is worth mentioning, that striatin was shown to be a regulator of vesicular trafficking in neurons (55).

### GUL1 interacts with the STRIPAK and MOR complexes

Our phosphoproteome results indicate that GUL1 is more highly phosphorylated in STRIPAK deletion mutants than in the wild type. The phosphorylation-dependent function of GUL1 is reminiscent of the findings for the yeast homolog SSD1. In yeast, nine predicted phosphorylation sites were functionally analyzed by mutagenesis. The phospho-deficient SSD1^S/T9A^ protein, where all nine sites were mutated, localizes to P-bodies and bound mRNAs disintegrate. However, this strain is only viable when an inducible promoter is used for gene expression. In contrast, the phospho-mimetic SSD1 variant SSD1^S/T9D^ is viable under constitutive gene expression and shows a polarized localization similar to the wild type protein (22). SSD1 is further involved in the regulation of translation of proteins involved in cell wall remodeling (20, 21), and its activity is dependent on the state of phosphorylation, which is determined by the NDR kinase Cbk1p, which interacts physically with SSD1 (22, 23, 37).

Our functional investigation of phospho-deficient and mimetic mutants also demonstrates that phosphorylation of GUL1 at S180 and S216 is critical for vegetative growth. S180 from GUL1 corresponds to the phosphorylation site S164 in SSD1, while the sites corresponding to GUL1 S216 and S1343 are not predicted as phosphorylation sites in the yeast protein. Moreover, the phosphorylation of GUL1 seems to be dependent on different signaling complexes, as proposed in our new model depicted in Fig 8. While S180 has a conserved recognition site for the NDR kinase, namely COT1, S216 is most probably phosphorylated by a casein kinase. From our phosphoproteome data, it therefore follows that S180 is dephosphorylated by STRIPAK, while a yet unknown phosphatase acts on S216. COT1, which was intensively investigated in *N. crassa*, is part of the MOR complex, and is regulated by the upstream GCK POD6. All components of the MOR complex are crucial for the polar organization of the actin cytoskeleton, and hence, fungal morphology (9, 16, 17). In *N. crassa, gul-1* deletion is able to partially suppress the phenotype of *cot-1*, and thus; is a dominant modifier of the NDR kinase COT-1, the homolog of the yeast kinase Cbk1p (14, 16, 17).

**Fig 8.**
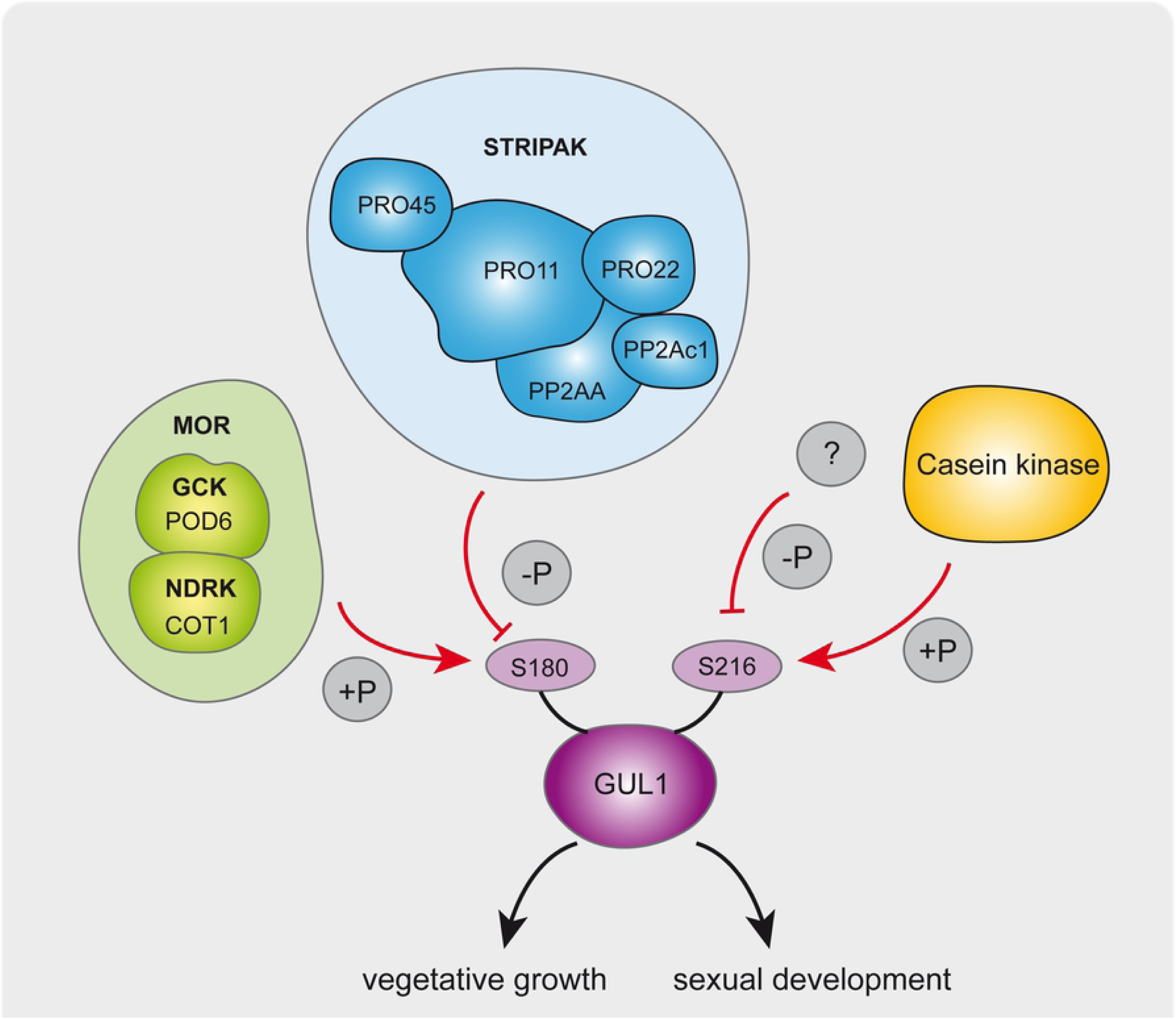
Schematic overview of phosphorylation dependent GUL1 function in sexual and asexual development. S180 is a molecular switch for hyphal growth and morphology, which is targeted by COT1 and STRIPAK. In contrast, S216 is probably targeted by casein kinase and a so far unknown phosphatase. Abbreviations: **MOR**= <u>m</u>orphogenesis <u>or</u>b6 network; STRIPAK: <u>s</u>triatin-interacting <u>p</u>hosphatase <u>a</u>nd <u>k</u>inase; **GCK** = germinal centre kinase; **NDRK** = nuclear dbf2-related kinase

Taken together, both the global proteome and phosphoproteome analyses of three STRIPAK mutants reveal that GUL1, an RNA-binding protein, is a dephosphorylation target of STRIPAK, which most probably acts parallel of MOR. The function of GUL1 is phosphorylation dependent and it is involved in hyphal morphology and sexual development. This work thus contributes further to the notion that coordinated cellular development is feasible through the interplay of several cellular signaling pathways, including the STRIPAK signaling complex. Importantly, the identification of STRIPAK targets in this work will promote new studies in other organisms than fungi, which are of interest as regards identifying phosphorylation targets of the STRIPAK signaling complex.

## Materials and Methods

### Strains and growth conditions

Electro-competent *E. coli* XL1-Blue MRF’ cells (56) were used for the generation of recombinant plasmids. Chemical competent NEB5α-cells (NEB biolabs) were used for propagation of plasmid DNA after Q5-mutagenesis. The resulting strains were grown under standard laboratory conditions (57) and were selected by ampicillin resistance. *S. cerevisiae* strain PJ69-4A was used for construction of plasmids p07544_OEC and pDS23-gul1-DsRed by homologous recombination as described previously (58, 59). The yeast cells were grown according to standard protocols (60), and transformants were selected by screening for uracil prototrophy.

*S. macrospora* strains, as listed in S3 Table, were grown under standard conditions and transformed with recombinant plasmids as described before (61, 62). The transformants were selected on medium supplemented with either nourseothricin (50 mg/ml) or hygromycin B (80 U/ml) or both. Isolation of gDNA was performed as reported previously (61). Integration of wildtype and mutated genes was verified by PCR and sequencing (Eurofins Scientific, Ebersberg, Germany). To obtain homokaryotic strains, transformants were crossed and ascospores were isolated from recombinant fruiting bodies. Growth tests were performed with three biological replicates with three technical replicates each. Strains were inoculated in petri dishes with 20 ml of SWG agar medium as an 8-mm-diameter agar plug of the respective strain. Growth fronts were measured after 24 h and 48 h.

### Protein extraction, enrichment, and fractionation

Samples were prepared as recently described (11). A bicinchoninic acid assay (Pierce BCA protein concentration assay kit) was performed according to the manufacturer’s protocol to determine the protein concentration in the lysates. Free cysteine residues were then reduced by addition of dithiotreitol (DTT) to the samples to a final concentration of 10 mM and incubation for 30 min at 56°C. For subsequent alkylation, iodoacetamide (IAA) was added at a concentration of 30 mM and after incubation for 30 min at room temperature in the dark, excess of IAA was quenched by addition of 10 mM DTT. Samples were further purified by ethanol precipitation, and prior to digestion, precipitated pellets were resuspended in 40 µl of 6 M guanidinium hydrochloride (GuHCl). A final concentration of 0.2 M GuHCl was reached by addition of ammonium bicarbonate buffer (pH 7.8) and CaCl_2_ was added at a final concentration of 2 mM. After addition of trypsin at a 1:20 ratio (protease:substrate, w/w), samples were digested at 37°C for 14 h and digestion was stopped by addition of 10 % trifluoroacetic acid (TFA). Following a desalting step, peptides were quality controlled as described before (63) and dried completely using a SpeedVac. After resuspension in 0.5 M triethylammonium bicarbonate (pH 8.5), 150 µg of tryptic peptides per sample were labelled with iTRAQ 8-plex reagents (AB Sciex, Darmstadt, Germany) according to the manufacturer’s protocol. Samples were pooled and quenched and a 70 µg aliquot was taken for global proteome analysis. Thereof, 35 µg were subjected to fractionation by high-pH reversed phase liquid chromatography (RPLC) using an Ultimate 3000 HPLC (high performance liquid chromatography) (Thermo Scientific, Dreieich, Germany) equipped with a C18 column (BioBasic 18, 5 µm particle size, 300 Å pore size, 150 × 0.5 mm). Fraction collection was performed in concatenated mode with 1 min windows and a total of 20 fractions were collected.

The remaining multiplexed sample (1,130 µg) was dried under vacuum and subjected to phosphopeptide enrichment. A protocol described by (64) using titanium dioxide (TiO_2_, Titansphere TiO, 5 µm particle size, GL Sciences Inc, Japan) was used and adapted as described in (65). Enriched phosphopeptides were further fractionated by means of hydrophilic interaction liquid chromatography (HILIC) using an Ultimate 3000 HPLC (Thermo Scientific, Dreieich, Germany) equipped with a TSKgel Amide-80 column (250 μm × 15 cm, 2 µm particle size, Tosoh Bioscience, Japan) and 23 fractions were collected.

### LC-MS/MS analysis

All global- and phosphoproteome fractions were subjected to LC-MS/MS analysis using an Ultimate 3000 nanoRSLC HPLC coupled to a Q Exactive HF mass spectrometer (both Thermo Scientific, Bremen, Germany). For preconcentration, samples were loaded onto a precolumn (Pepmap RSLC, Thermo Scientific, C18, 100 µm x 2 cm, 5 μm particle size, 100 Å pore size) for 5 min at a flow rate of 20 µl/min (0.1 % TFA). Peptide separation on the analytical column (Pepmap RSLC, Thermo Scientific, C18, 75 μm x 50 cm, 2 μm particle size, 100 Å pore size) was performed at a flow rate of 250 nL/min. A binary gradient of solvent A (0.1 % formic acid (FA) and B (84 % acetonitrile, 0.1 % FA) was used with a linear increase of solvent B from 3 to 35 % in 120 min for global proteome fractions and 3 to 42 % in 100 min for phosphoproteome fractions. MS analysis was performed in a data-dependent acquisition (DDA) mode after first performing a survey scan from 300 to 1,500 m/z at a resolution of 60,000 and with the AGC target value set at 1 × 10^6^ and a maximum injection time of 120 ms. The top 15 most abundant precursor ions of every survey were selected for fragmentation by higher-energy collisional dissociation (HCD) and MS/MS analysis, and were dynamically excluded from selection for the following 30 s. MS/MS scans were acquired at a resolution of 15,000 and with the AGC target value set to 2 × 10^5^, a maximum injection time of 250 ms, and a fixed first mass of 90 m/z. For global proteome fractions, quadrupole precursor selection was performed with an isolation window width of 0.7 m/z and normalized collision energy (nCE) of 31 %, while for phosphoproteome fractions, precursors were isolated with an isolation window width of 1.0 m/z and fragmented with 33 % nCE. The polysiloxane ion at m/z 371.101236 was used as lock mass and a 10 % (v/v) NH_4_OH solution was placed at the nano source as described previously (66) to reduce precursor charge states.

### Proteomics data analysis

MS raw files were analyzed with Proteome Discoverer 1.4 (Thermo Scientific, Bremen, Germany) using the search algorithms Mascot (version 2.4.1, Matrix Science), Sequest HT, and MS Amanda. Searches were performed in target/decoy mode against a *S. macrospora* protein sequence database (10,091 target sequences) with the following parameters. Enzyme specificity was set to “trypsin”, allowing for a maximum of 2 missed cleavages. Precursor mass tolerance was limited to 10 ppm and fragment mass tolerance to 0.02 Da. iTRAQ 8-plex at peptide N-termini and lysine residues as well as carbamidomethylation of cysteines were set as fixed modifications. Oxidation of methionine was allowed as a variable modification in all searches and phosphorylation of serine, threonine, or tyrosine was additionally set as a variable modification for phosphoproteome analysis. To determine the modification site confidence in the latter case, phosphoRS node (version 3.1; (67) was used (S6 Fig). False discovery rate (FDR) estimation was performed by the Percolator node (68) and results were filtered to 1 % FDR on the peptide spectrum matches (PSM) level, only allowing for rank 1 hits. A minimum of 2 unique peptides per protein were required for global proteome data and only phosphorylated peptides with a phosphoRS site probability ≥ 90 % were exported for phosphoproteome analysis. Global proteome data was normalized to correct for systematic errors during sample labelling by implementation of correction factors based on the summed total intensities of all iTRAQ channels. After which, mean protein abundances of all biological replicates were calculated and ratios of the knockout strains against the wildtype were determined and log2 transformed. Only proteins exhibiting an absolute log2 ratio greater than two times the standard deviation of all proteins of the respective condition were considered as regulated. An Excel macro provided by (67) was used for analysis of phosphoproteome data. The correction factors determined from the global proteome data was used for normalization and only ratios of confidently localized phosphorylations were used. Ratios were calculated as described above and only phosphopeptides exhibiting an absolute log2 ratio greater than two times the standard deviation of the respective proteins in the global proteome data were considered as regulated.

### Phosphorylation motif analysis

To identify overrepresented consensus motifs of the identified phosphorylation sites, seven flanking amino acids up- and downstream of the modified residues were extracted. The motifs of up- or downregulated sites in the individual deletion strains were uploaded to the MoMo web server (69). Significantly enriched motifs were identified using the motif-x algorithm and the *S. macrospora* protein database (10,091 sequences) as context sequence and requiring a minimum number of 20 occurrences and a p-value of threshold of 1E^-6^.

### Generation of deletion strains

To generate a Δ*gul1* strain, a circular pKO-gul1 plasmid was transformed into a Δ*ku70* strain (70), and primary transformants were selected for hygromycin B resistance. Ascospore isolates of the Δ*gul1* strain with the genetic background of the wildtype were obtained as described before by crosses against the spore color mutant fus (32, 61) and verified by resistance to hygromycin B and sensitivity to nourseothricin. To obtain a *gul1pro45* double-deletion strain, Δ*pro45* (24) with a wildtype genetic background was crossed against *Δgul1/fus*. Ascospores from tetrads were selected for their hygromycin B resistance. All strains were verified by PCR and Southern blot analyses (S7 Fig and S8 Fig). The Δgul1 strain was complemented using p07544_OEC, which encodes a *gul1-gfp* fusion gene under the control of the constitutive *gpd* promotor. Phospho-mutants were generated by transformation of the mutated plasmids (S4 Table) into the Δgul1 strain. Phospho-mutations in the generated strains were verified by PCR analysis and DNA sequencing (Eurofins Genomics; Ebersberg, Germany). The expression of the mutated genes was verified by a Western blot analysis (S4 Fig). Unless otherwise stated, all wildtype and mutant strains carry the *fus* mutation, which results in reddish ascospores (32).

### *In vitro* recombinant techniques and construction of phospho-mutants

Plasmid constructions were performed via either conventional restriction and ligation with T4 DNA ligase or homologous recombination in yeast (59). For phospho-mimetic and -deficient strains, plasmid p07544_OEC carrying *gul1* was used for Q5 mutagenesis (NEB biolabs). Using specific primers (S5 Table), we generated four plasmids, containing phospho-mimetic and phospho-deficient mutations (S9 Fig). After DNA-mediated transformation of the abovementioned plasmids into Δ*gul1*, we obtained homokaryotic single spore isolates of phospho-mimetic strains S180E, S216E, and S1343E and of the phospho-deficient strains S216A and S1343A. However, we failed in generating homokaryotic isolates of the phospho-deficient strain S180A. In total, we investigated 340 ascospores from two independent primary transformants. From 105 germinated ascospores, none showed resistance against nourseothricin, indicating that the ascospores do not carry the *gul1*-complementation vector. This result strongly suggests that the phospho-deficient mutation S180A is lethal, and only heterokaryotic strains are selected on nourseothricin-containing plates. For our further analysis, we investigated a primary transformant S180A that is considered to be heterokaryotic.

### Microscopic investigations

Microscopic investigations were performed with an AxioImager microscope (Zeiss, Jena, Germany). Sexual development was documented by differential interference contrast (DIC) microscopy with strains inoculated on BMM-coated glass slides in petri dishes for 7 to 10 days. To analyze ascus rosettes, mature perithecia were isolated and opened mechanically. To analyze septation and hyphal morphology, strains were grown on minimal-starch-medium (MMS)-coated glass slides in petri dishes for 2 days. (61, 71). Co-localization of proteins was carried out by inoculation of two different strains on the same BMM-coated glass slides in petri dishes for 1 to 2 days. Hyphal fusion of both strains enabled the formation of heterokaryons by exchanging nuclei. Microscopic investigations were carried out with an AxioImager M.1 microscope (Zeiss) equipped with a CoolSnap HQ camera (Roper Scientific) and a SpectraX LED lamp (Lumencor). GFP, mRFP, and DsRed fluorescence were analyzed using filter set (Chroma Technology Corp.) 49002 (GFP, excitation filter HQ470/40, emission filter HQ525/50, beamsplitter T495LPXR) or 49008 (mRFP and DsRed, excitation filter HQ560/40, emission filter ET630/75m, beamsplitter T585lp). Calcofluor White M2R (CFW) fluorescence was analyzed using Chroma filter set 31000v2 (excitation filter D350/50, emission filter D460/50, beam splitter 400dclp; Chroma Technology Corp., Bellows Falls, VT, USA). For fluorescence microscopy, strains were grown on BMM-coated glass slides for 1 to 2 days (61). For analysis of directed movement images were captures with an Orca Flash4.0 camera (Hamamatsu, Japan) and objective lens Plan Apochromat (63x, NA 1.4). Fluorescently-labeled proteins were excited using a laser-based epifluorescence-microscopy. A VS-LMS4 Laser Merge-System (Visitron Systems) combines solid state lasers for the excitation of Gfp (488 nm/100 mW) and Rfp/mCherry (561 nm/150 mW). All parts of the microscope systems were controlled by the software package VisiView (Visitron). Kymographs were generated as described previously (72). Staining with Calcofluor White M2R (Sigma-Aldrich) was performed with a 1 μg/ml CFW stock solution diluted 1:400 in a 0.7% NaCl solution. Staining with FM4-64 (Invitrogen) was performed with a concentration of 5 µg/ml and incubation of 1 min on ice. Images were captured with a Photometrix Cool SnapHQ camera (Roper Scientific) and MetaMorph (version 6.3.1; Universal Imaging), and further processed with MetaMorph and Adobe Photoshop CS6. Videos were processed with Adobe Media Encoder CS6 (Adobe Systems Inc.). The time scale for the videos corresponds to seconds. Quantification of perithecia was obtained by counting mature perithecia under a binocular (Zeiss) within 1 cm^2^. These experiments was performed for three biological replicates with three technical replicates each.

### Data availability

The mass spectrometry proteomics data were deposited to the ProteomeXchange Consortium via the PRIDE partner repository (73) with the dataset identifier PXD016296.

## Acknowledgements

We thank Ingeborg Godehardt and Susanne Schlewinski for superb technical help, Dr. Daria Radchenko for construction of double mutant Δpro45Δpro11, Ramona Lütkenhaus for providing strain H2A-mRFP/fus and Prof. Dr. S. Pöggeler (Göttingen) for providing *S. macrospora* strains expressing *rab5*- and *rab7.*

## Author Contributions

Conceptualization: Valentina Stein, Ulrich Kück

Data curation: Valentina Stein, Bernhard Blank-Landeshammer, Kira Müntjes

Formal analysis: Valentina Stein, Bernhard Blank-Landeshammer, Kira Müntjes, Ramona Märker

Funding acquisition: Ines Teichert, Michael Feldbrügge, Albert Sickmann, Ulrich Kück

Investigation: Valentina Stein, Bernhard Blank-Landeshammer, Kira Müntjes, Ramona Märker

Methodology: Valentina Stein, Bernhard Blank-Landeshammer, Kira Müntjes, Ramona Märker

Project administration: Ulrich Kück

Resources: Ines Teichert, Michael Feldbrügge, Albert Sickmann, Ulrich Kück Software: Valentina Stein, Bernhard Blank-Landeshammer

Supervision: Michael Feldbrügge, Albert Sickmann, Ulrich Kück Validation: Albert Sickmann, Ulrich Kück

Visualization: Ulrich Kück

Writing – original draft: Valentina Stein, Bernhard Blank-Landeshammer, Ines Teichert, Michael Feldbrügge, Albert Sickmann, Ulrich Kück

Writing – review & editing: Valentina Stein and Ulrich Kück

## Supporting information

**S1 Fig. Proteins identified and quantified in this and the previous study** (11). (A) In total 4,349 proteins were quantified in this study, compared to 4,193 in our previous study, 93 % of which we were covered in this study. (B) The commonly used deletion strain Δpro11 was used to compare the quantification between the two analyses and a Pearson’s correlation coefficient of 0.7339 was calculated.

**S2 Fig. Phosphoproteins and –peptides identified and quantified in this and the previous study (11).** (A, C) In total 9,773 phosphopeptides originating from 2,465 proteins were quantified in this study, compared to 10,635 phosphopeptides from 2,489 phosphoproteins in the previous study (11), 58 % and 84 % of which were commonly identified, respectively. (B) The deletion strain Δpro11 was used to compare the quantification between the two analyses and a Pearson’s correlation coefficient of 0.621 was calculated for the commonly identified phosphopeptides.

**S3 Fig. Phenotype of Δgul1 hyphae in different regions of the colony.** Strains were grown on MMS and cellophane for four days. Wild type served as control. Dotted lines indicate the hyphal area of microscopic images. Not drawn to scale.

**S4 Fig. Expression control of phospho-mutated variants of GUL1 tagged with GFP.** Strains were grown for 3 days in liquid media (BMM) as a surface culture. For each strain, 10 µg of crude protein extract were subjected to SDS-PAGE. Western blot analysis was performed with an anti-GFP antibody and an anti-α-Tubulin antibody as control. GUL1 tagged with GFP has a mass of 175 kDa, while α-Tubulin has a mass of 55 kDa. GUL1-GFP was detected in all six different phospho-mutants (S180A and S180E, S216A, S216E, S1343A and S1343E). Wild type and a complemented Δgul1 strain were used as control.

**S5 Fig. Shuttling signals of GUL1-GFP phospho-variants.** Examples of kymographs, used for the analysis of moving GUL1. For this analysis, kymographs were generated for a distance of 50 µm 20 µm beyond the hyphal tip. The shuttling of GUL1-GFP was measured in 20 different hyphae per strain.

**S6 Fig. Example for a tandem mass spectrum of the peptide TRSDSKVPVGDTPEAR, identifying phosphorylation of GUL1 residue S1289.** Y-ions are depicted in blue, b-ions in red, b-ions with neutral loss of H_3_PO_4_ in pink and iTRAQ reporter ions in purple. B- and y ions were used for scoring by the Mascot search algorithm, while all ions were used by the phosphoRS algorithm (67) to calculate the phosphorylation site probability of 99.6 % for this peptide The b_3_-P, b_4_-P and and y_12_ ions are indicative of the phosphorylation on serine 3.

**S7 Fig. Deletion strategy and verification of a *gul1* deletion strain at the *gul1* locus via PCR and Southern blot analysis.** (A) Genomic situation of the wild type and Δgul1. Genes are indicated by arrows showing primers for the verification of the deletion via PCR fragments, which are shown as grey lines. The thick grey line indicate DNA fragments used as probes for Southern hybridization. The restriction sites of the enzyme *Hin*dIII are displayed, which was used for restriction of the DNA for Southern blot analysis. Dotted lines display areas for homologous integration. Not drawn to scale (B) PCR analysis for the verification of the *gul1* deletion. Integration of 5’-flank *gul1*, 3’-flank *gul1* and *gul1* was tested. Genomic DNA of the wild type (wt) served as control. Negative control (NK) contained no DNA. (C) Autoradiograph of Southern blot hybridization with radioactively labeled probes specific for *gul1* and *hph* after digestion of the genomic DNA of wt and the *gul1* deletion strain with *Hin*dIII.

**S8 Fig. Deletion strategy and verification of double deletion of *gul1* and *pro45* via PCR and Southern blot analysis.** (A) Genomic situation of the wt, Δgul1 and Δpro45. Arrows indicate primers for the verification of the deletion via PCR, which are shown as black lines. The thick grey lines indicate DNA fragments used as probe for Southern hybridization. The restriction sites of the enzymes are displayed, which were used for restriction of the DNA for Southern blot analysis. Dotted lines display areas for homologous integration. (B) PCR analysis for the verification of the *gul1*- and *pro45* deletion. Integration of 5’-flank *gul1*, 3’-flank *gul1* and *gul1* was tested, as well as 5’-flank *pro45*, 3’-flank *pro45* and *pro45* in S156228. Genomic DNA of the wt served as control. Negative control (NK) contained no DNA. (C) Autoradiograph of Southern blot hybridization with radioactively labeled probes specific for *hph, gul1* and *pro45.* Genomic DNA for hybridization with *gul1, pro45 and hph* was digested with *Hin*dIII, *Eco*RI and *Pvu*II, respectively.

**S9 Fig. Phospho-mimetic and –deficient versions of *gul1* used for functional analysis**. Lowercase and capital letters indicate the coding sequence of *gul1* and the derived amino acid sequence, respectively, close to serine phosphorylation sites S180, S216 and S1343. The triplets encoding the phosphorylated amino acids are given in bold letters and highlighted in grey. Red letters indicate single base pair substitutions and the corresponding amino acid substitutions S180A, S180E, S216A, S216E. S1343A and S1343E.

**S1 Table. Identified phosphorylation sites of GUL1 in (11).** The phosphoproteomic study of Δpp2Ac1, Δpro11, and Δpro22 compared to the wild type identified seven phosphorylation sites in GUL1. None is differentetially phosphorylated in the three STRIPAK single deletion strains. For each phosphorylation site of GUL1, log2 ratio of reporter ion intensity in deletion strain and wild type relative to the respective standard deviation is given. Bold numbers indicate an upregulation of the phosphorylation site compared to the wild type. Regular numbers indicate no regulation of the phosphorylation site compared to the wild type. Standard deviations of the ratio of phosphopeptides in mutants compared to wild type: Δpp2Ac1: 0.63; Δpro11: 0.61; pro22: 0.49.

**S2 Table.** Values for vegetative growth rates of single and double mutant strains to evaluate genetic interactions.

**S3 Table. Strains used in this work**

**S4 Table. Plasmids used in this work**

**S5 Table. Oligonucleotides used in this work**

**S1 Movie. Shuttling of GUL1-GFP.** GUL1-GFP shuttles like PAB1-positive endosomes in hyphae (20 µm beyond hypal tip shown, hyphal tip towards the right; scale bar, 50 µm; timescale in seconds, 150 ms exposure time, 150 frames, 6 frames/s display rate, MPEG-4 format; corresponds to Fig 7A).

**S2 Movie. Shuttling of GFP-RAB5.** GFP-RAB5 shuttles like PAB1-positive endosomes in hyphae (20 µm beyond hypal tip shown, hyphal tip towards the right; scale bar, 50 µm; timescale in seconds, 150 ms exposure time, 150 frames, 6 frames/s display rate, MPEG-4 format; corresponds to Fig 7A).

**S3 Movie. GFP-RAB7.** GFP-RAB7 in hyphae (20 µm beyond hypal tip shown, hyphal tip towards the right; scale bar, 50 µm; timescale in seconds, 150 ms exposure time, 150 frames, 6 frames/s display rate, MPEG-4 format; corresponds to Fig 7A).

**S4 Movie. Co-localization of GUL1-GFP and GFP-PAB1.** Processive GUL1-DsRed signals co-migrate with GFP-PAB1 in hyphae (20 µm beyond hyphal tip, hyphal tip towards the right; scale bar, 50 µm; timescale in seconds, 150 ms exposure time, 150 frames, x6 frames/s display rate; MPEG-4 format; corresponds to Fig 7C).

**S1 Dataset. Sordaria_iTRAQ_pH8_Proteins S2 Dataset. iTRAQ8Plex_Phospho_Sordaria**

